# Bidirectional regulation of glycoprotein nonmetastatic melanoma protein B by β-glucocerebrosidase deficiency in *GBA1* isogenic dopaminergic neurons from a patient with Gaucher disease and parkinsonism

**DOI:** 10.1101/2025.06.23.661126

**Authors:** Chase Chen, Charis Ma, Richard Sam, Jens Lichtenberg, Tiffany Chen, Ying Hao, Ziyi Li, Isabelle Kowal, Kate Andersh, Yue Andy Qi, Gani Perez, Ellen Hertz, Yan Li, Darian Williams, Mark J. Henderson, Morgan Park, Xuntian Jiang, Pilar Alvarez Jerez, Cornelis Blauwendraat, Ellen Sidransky, Yu Chen

**Affiliations:** Molecular Neurogenetics Section, Medical Genetics Branch, National Human Genome Research Institute, National Institutes of Health, Bethesda, Maryland, USA; Aligning Science Across Parkinson’s (ASAP) Collaborative Research Network, Chevy Chase, MD, 20815; Center for Alzheimer’s and Related Dementias (CARD), National Institute on Aging and National Institute of Neurological Disorders and Stroke, National Institutes of Health, Bethesda, MD, USA; DataTecnica LLC, Washington DC, USA; Proteins/Peptides Sequencing Facility, National Institute of Neurological Disorders and Stroke, National Institutes of Health, Bethesda, Maryland, USA; Division of Preclinical Innovation, National Center for Advancing Translational Sciences, National Institutes of Health, Rockville, Maryland, USA; NIH Intramural Sequencing Center, National Human Genome Research Institute, National Institutes of Health, Rockville, Maryland, USA; Diabetic Cardiovascular Disease Center, Washington University School of Medicine, St. Louis, Missouri, USA; Laboratory of Neurogenetics, National Institute on Aging, National Institutes of Health, Bethesda, Maryland, USA

**Keywords:** *GBA1*, GCase, Gaucher disease, Parkinson disease, GPNMB, iPSC

## Abstract

Variants in *GBA1* are common genetic risk factors for several synucleinopathies. The increased risk has been attributed to the toxic effects of misfolded glucocerebrosidase (GCase) (gain-of-function), and the accumulation of lipid substrates due to reduced enzyme activity (loss-of-function). To delineate *GBA1* pathogenicity, an iPSC line was generated from a patient with both type 1 Gaucher disease (*GBA1*: N370S/N370S; p.N409S/p.N409S) and Parkinson disease (PD). From this line, we created a reverted wild-type (WT) line and a *GBA1* knockout (KO) line to eliminate misfolded GCase and intensify lipid accumulation. N370S/N370S and KO dopaminergic neurons (DANs) exhibited decreasing GCase levels and progressive accumulation of lipid substrates compared to WT DANs. Notably, the expression of *GPNMB*, whose levels correlate with PD risk, was upregulated by the mild lipid accumulation in N370S/N370S DANs but disrupted in KO DANs. These findings refine the loss-of-function mechanism by associating PD risk levels of GPNMB with lipid levels specific to *GBA1* risk variants.

## Introduction

Synucleinopathies, including Parkinson disease (PD) and dementia with Lewy bodies (DLB), are complex movement disorders resulting from a combination of aging, environmental exposures, and genetic factors^1^. Our understanding of the genetic basis of PD has advanced considerably, with linkage studies in large families with PD revealing highly penetrant rare variants^2,3^ and genome-wide association studies (GWASs) identifying more common, smaller effect variants in apparently sporadic PD^4–6^.

*GBA1* variants are the most common genetic risk factor for PD^7,8^. *GBA1* encodes the lysosomal hydrolase β-glucocerebrosidase (GCase), which hydrolyses the glycosphingolipids glucosylceramide (GlcCer) and glucosylsphingosine (GlcSph)^9^. Biallelic pathogenic *GBA1* variants lead to Gaucher disease (GD), a rare lysosomal storage disorder^10^. The observation of parkinsonism in a rare subgroup of individuals with GD first directed attention to the role of GCase deficiency in the pathogenesis of PD^11,12^. PD also occurs more frequently in people heterozygous for Gaucher *GBA1* variants, and 3–25% of people with PD carry a Gaucher *GBA1* variant^7,8^. Additionally, PD risk is also associated with non-Gaucher causing *GBA1* variants, including the E326K (p.E365K) variant prevalent in PD cases in UK^13^ and the African ancestry-specific variant (rs3115534-G) present in ∼50% of West African individuals with PD^14^. However, only a small percentage of individuals with *GBA1* variants develop parkinsonism, indicating that the penetrance is low^15^ and implicating the contribution of genetic modifiers^16^. PD risk associated with *GBA1* variants has been attributed to the toxic effects of misfolded glucocerebrosidase (GCase) (gain-of-function)^17–20^, and/or the accumulation of lipid substrates due to reduced enzyme activity (loss-of-function)^21–23^. Additionally, *GBA1* has a highly homologous pseudogene *GBAP1* that is expressed at the mRNA and protein levels with functions yet to be defined^24^.

*GPNMB*, encoding the transmembrane protein glycoprotein nonmetastatic melanoma protein B (GPNMB), is a PD risk gene identified by GWAS^25^, and the PD risk–associated haplotype is associated with an approximately three-fold higher *GPNMB* expression in multiple brain regions^25,26^. Upregulated GPNMB levels have been proposed to increase PD risk by promoting the internalization of fibrillar α-synuclein and subsequent development of α-synuclein pathology^26^. GPNMB and its homolog PMEL17, first identified as melanosomal proteins, share a similar domain structure encompassing a large luminal domain containing a N-terminal domain (NTR), a polycystic kidney disease-like domain (PKD) and a kringle-like domain (KRG), and an intracellular domain, also referred to as the C-terminal domain (CTD), with a dileucine-based lysosomal sorting signal^27,28^. PMEL17 is processed by a proprotein convertase in the Golgi complex^29^ and by a metalloprotease of the ADAM family in the melanosomes^30^ to produce fragments containing the PKD and the repeat (RPT) domains, which eventually assemble into mature fibrils to facilitate the polymerization of melanins^31^.

In this work, we sought to gain a deeper understanding of the biologic consequences of *GBA1* variants by probing GCase-associated cellular and lysosomal endpoints in high-purity dopaminergic neurons (DANs) differentiated from *GBA1* isogenic iPSC lines. These unique GD/PD patient iPSC lines enabled a decoupling of the consequences of GCase variants, namely misfolded GCase versus lipid accumulation, and presented distinct lipid substrate levels. Remarkably, GPNMB levels were elevated in N370S/N370S DANs but disrupted in knockout (KO) DANs, revealing a dosage-dependent regulation of GPNMB by lipid storage and implicating GPNMB in *GBA1* pathogenicity.

## Results

### Generation of iPSC line HT809 from a patient with both Gaucher disease and Parkinson disease

The proband was an Ashkenazi Jewish male diagnosed with type 1 GD (*GBA1*: N370S/N370S) at age 61 (Fig. S1A), when he presented with thrombocytopenia, osteopenia, and mild splenomegaly. He was not treated with enzyme replacement therapy. At age 64 he developed a tremor of the right hand, which progressed and was accompanied by rigidity, bradykinesia, stiffness, shuffling gait, and impaired olfaction. His cognition, evaluated at ages 65, 67 and 74, was reported as normal. He died at age 76 from complications of cardiac surgery^32^. A skin biopsy was performed at age 65, after the development of parkinsonism, and skin fibroblasts were reprogrammed to the iPSC line HT809 using CytoTune-iPS 2.0 Sendai Reprogramming Kit (Fig. S1B,C).

To gain insight into the patient’s genetic background, we sequenced his genome at >30x coverage. The distribution of insertion-deletion (indel), loss-of-function (LoF), and missense SNVs was similar to human populations reported in the gnomAD database, as well as the recently established reference iPSC line KOLF2.1J (Fig. 1A)^33^. After restricting our analysis to variants with a gnomAD allele frequency (AF) of <0.001 and a combined annotation-dependent depletion (CADD) phred score of >30, a modest number of potentially deleterious variants were found (Fig. S3A N370S/N370S), but none had an established association with PD, other than the N370S *GBA1* variant. Finally, we calculated polygenic risk scores (PRSs)^34^ for HT809 based on the cumulative burden of common genetic variants associated with Alzheimer disease (AD) or PD and found that his PRSs fell within the expected population range (Fig. 1B,C). Together, these findings demonstrate that HT809 is a representative case for GD/PD with a relatively neutral genetic background when *GBA1* variants are excluded. We refer to this parental N370S/N370S line as “HT809-N370S” in this study.

**Figure 1:**
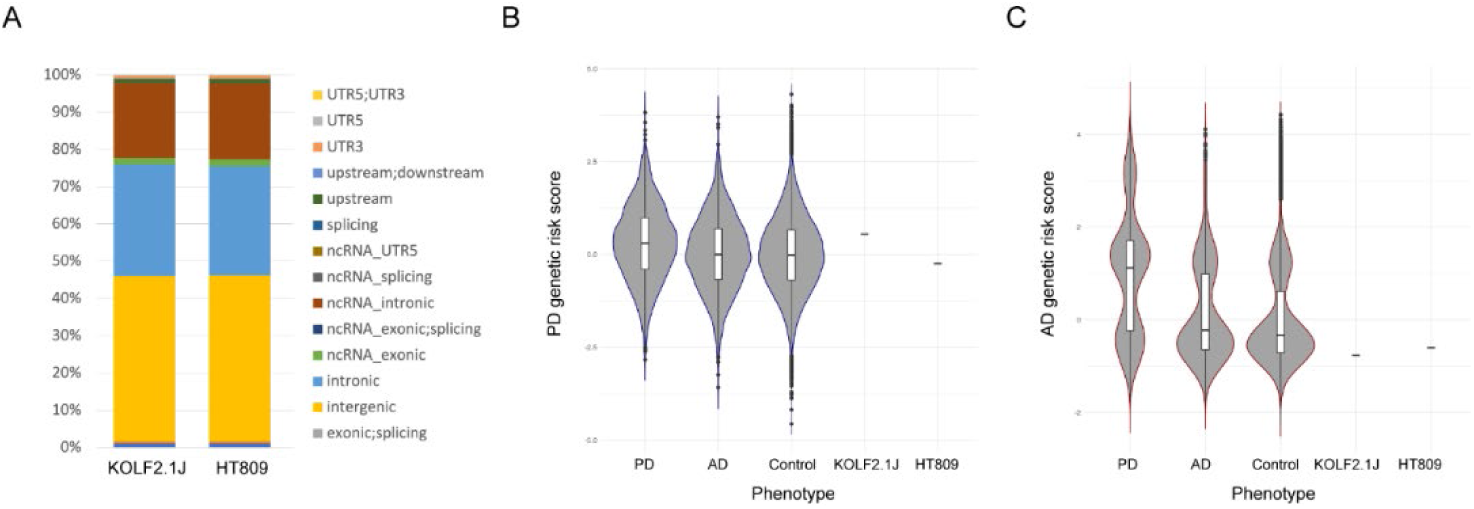
Genetic background of HT809. **A.** The distribution of insertion-deletion (indel), loss-of-function (LoF), and missense SNVs in HT809 and KOLF2.1J. **B.C.** PD and AD polygenic risk scores for HT809 and KOLF2.1J.

### Engineering HT809 to facilitate the evaluation of dopaminergic neuronal differentiation and to enable LysoIP

Before editing the *GBA1* alleles, we engineered HT809 to facilitate the monitoring of dopaminergic neuron (DAN) differentiation and to enable rapid purification of lysosomes through LysoIP. Tyrosine hydroxylase (TH), encoded by *TH*, is the rate-limiting enzyme involved in dopamine synthesis, and is a molecular marker for DANs. We inserted tdTomato, a red fluorescent protein, at the C-terminus of the *TH* gene locus (Fig. 2A)^35^. In the targeted locus, the *TH* gene stop codon was removed and a P2A self-cleaving peptide preceded the tdTomato sequence, placing the expression of tdTomato under the control of the endogenous *TH* promoter. When adapting the DAN differentiation protocol based on a two-step WNT signaling activation strategy^36^, we determined the optimal concentration of the glycogen synthase kinase 3 (GSK3)-inhibitor CHIR99021 for the “CHIR boost” in HT809 to be 3 µM, with higher concentrations (5 µM and 7 µM) triggering dose-dependent cell death and decreased yield of differentiated cells (Fig. 2B). On differentiation day 25, ∼75% of the cells expressed tdTomato, suggesting DAN identity (Fig. 2C), which was further confirmed by the typical neuronal morphology observed for FACS-sorted tdTomato+ cells (Fig. 2D). When co-stained for tdTomato and TH, TH was detected in all tdTomato+ DANs, establishing tdTomato as a reliable reporter for TH (Fig. 2D). FOXA2, another midbrain floor plate marker, was also expressed in tdTomato+ DANs (Fig. 2D). Gene expression analysis using RNA sequencing (RNA-seq), performed on day 75 of differentiation, revealed a clear increase in a panel of DAN makers (Fig. 2E) and an enrichment of gene sets involved in brain and neuron development (Fig. 2F).

**Figure 2:**
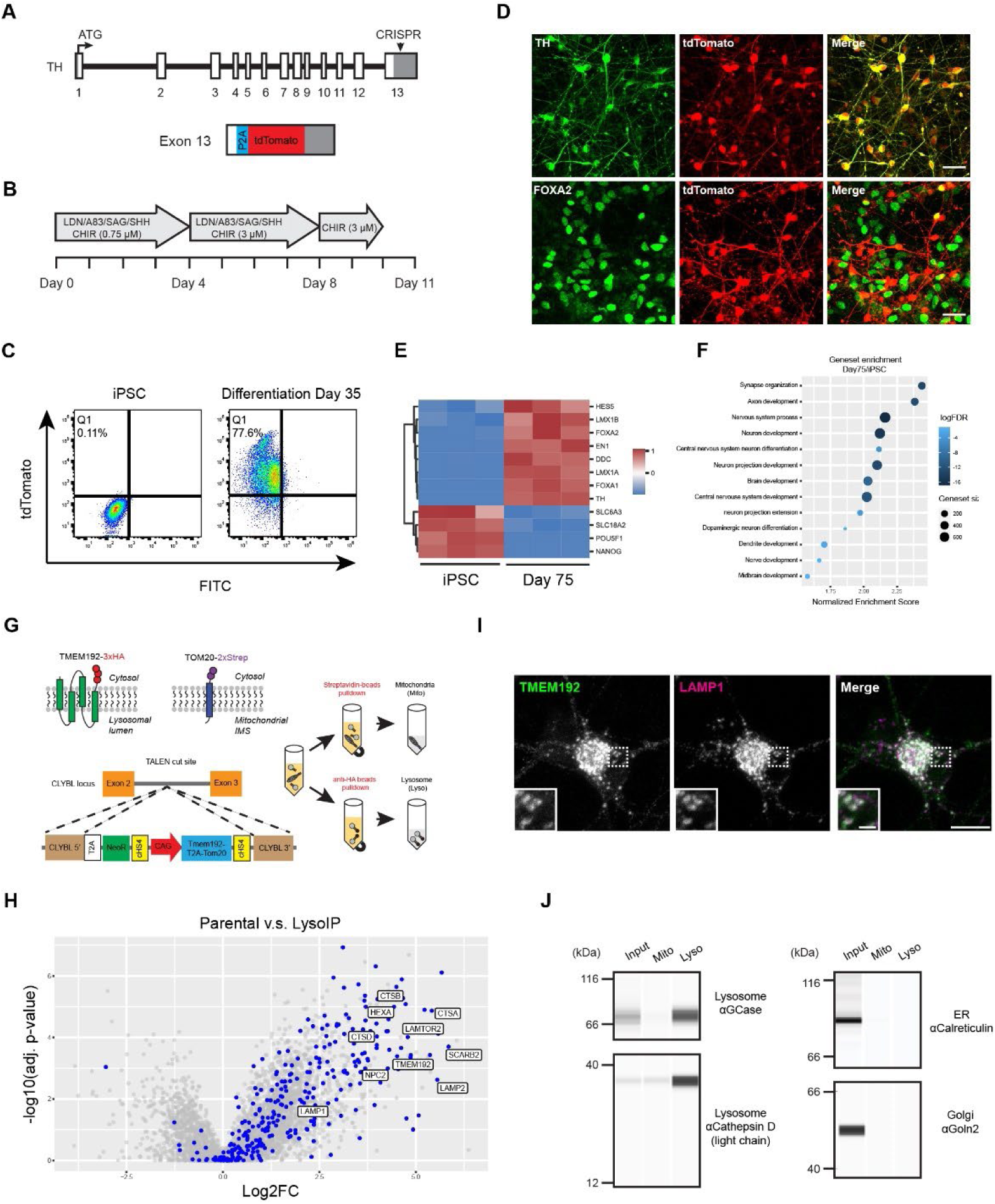
Engineering HT809 to monitor DAN differentiation and to enable LysoIP. **A.** Insertion of tdTomato into the endogenous *TH locus.* B. DAN differentiation protocol optimized for HT809. **C.** DAN differentiation efficiency evaluated by FACS. **D.** Immunostaining TH and FOXA2 in tdTomato+ cells. Scale bar, 35 µm. **E, F.** RNA-seq comparing HT809 iPSCs and day 75 DANs. **G.** LysoIP and MitoIP strategy in HT809. **H.** Enrichment of lysosomal proteins in LysoIP samples. Lysosomal proteins are indicated with blue color. **I.** Localization of TMEM192-GFP-3xHA to lysosomes in DANs. Scale bar, 10 µm; insert scale bar, 2 µm. J. Characterization of lysosomes purified from DANs by Western blotting using the Automated Western Blot JESS System.

To enable LysoIP^37^ and MitoIP^38^ in HT809 DANs, we inserted into the *CLYBL* safe harbor site^39^ a transgene cassette to express TMEM192-GFP-3xHA and TOM20-BFP-twinSTREP, separated by a T2A sequence. The transgene cassette was driven by a CAG promoter active in both iPSCs and differentiated neurons^39^. TMEM192-GFP-3xHA expression was higher than the endogenous TMEM192 and had a molecular weight of ∼66 kD, which is consistent with the sum of the molecular weights of TMEM192 (∼45 kD) and GFP (∼27 kD) (Fig. S2A). Additionally, TOM20-BFP-twinSTREP localized to mitochondria labeled with mitofusin, and TMEM192-GFP-3xHA to lysosomes labeled with LAMP1 (Fig. S2B). Together, these data indicate the efficient cleavage of the T2A sequence and the correct targeting of tagged TMEM192 and TOM20. As our priority was to characterize lysosomes, we adapted the LysoIP workflow in this study. Lysosomes purified from iPSCs showed enrichment of a large group of lysosomal proteins by proteomics analysis (Fig. 2H). TMEM192-GFP-3xHA expression and its lysosomal localization were maintained after DAN differentiation (Fig. 2I). Western blot analysis of lysosomes purified from DANs further confirmed the enrichment of lysosomal proteins including GCase and Cathepsin D, without contamination from other organelles such as the endoplasmic reticulum (ER) or the Golgi (Fig. 2J).

### Generation of HT809 isogenic lines harboring *GBA1* knockout and wild type

Located on chromosome 1q21, *GBA1* contains 11 exons and 10 introns over a length of 7.6 kb. 16 kb downstream of *GBA1* is the shorter (5.7 kb) pseudogene, *GBAP1* (Fig. 3A). Within coding regions, *GBA1* is 96% homologous to *GBAP1*, and this homology increases further to ∼98% in the region between intron 8 and the 3′ UTR. The scarcity of distinct features between *GBA1* and *GBAP1* sequences presents challenges to designing guide RNAs (gRNAs) to target *GBA1* while avoiding unintended editing of *GBAP1* that may perturb its newly discovered roles^24^. Aiming to knock out *GBA1* in HT809 by introducing frameshift indel mutations, we evaluated several gRNAs targeting *GBA1* exon 4. One gRNA, with four bases that differed between the *GBA1* target site and the potential *GBAP1* off-target site (Fig. 3B), demonstrated both editing efficiency and specificity. One iPSC clone edited with this gRNA acquired the desired frameshift mutations in both *GBA1* alleles with no change in *GBAP1* (Fig. 3C) and is referred to here as the *GBA1* homozygous knockout (KO) line HT809-KO.

**Figure 3:**
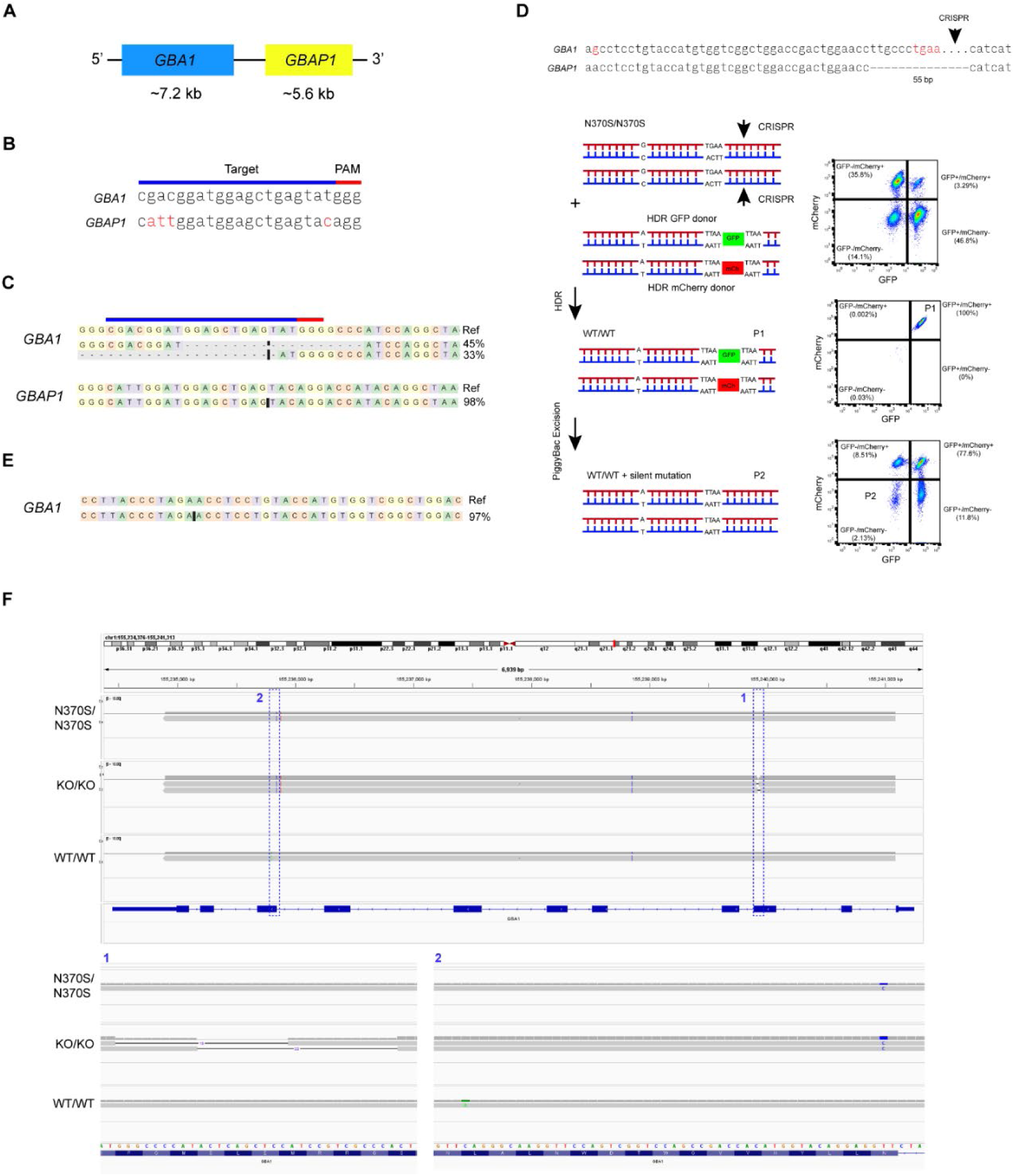
*GBA1* isogenic iPSC lines generated using CRISPR-Cas9 and piggyBac transposase. **A.** Schematic representation of *GBA1* and *GBAP1*. **B.** gRNA specifically targets *GBA1* to generate the KO line. **C.** Editing outcome in the KO line revealed by short-read amplicon sequencing. **D.** Reversion to the WT line using CRISPR-Cas9 and piggyBac transposase. **E.** Editing outcome in the WT line revealed by short-read amplicon sequencing. **F**. Specific editing of *GBA1* as demonstrated by PacBio HiFi long-read amplicon sequencing.

Homozygosity for the N370S variant in HT809 resulted from biallelic adenine (A) to guanine (G) changes in *GBA1* exon 10 (c.1226A>G). *GBA1* and *GBAP1* share identical sequence in the 61-bp region flanking the A-to-G mutation, undermining the feasibility of specifically targeting *GBA1* in this region. Since 39-bp downstream of the A-to-G point mutation *GBAP1* lacks a 55-bp sequence in exon 10, which is the defining exonic difference between *GBAP1* and *GBA1*^40^, we specifically targeted *GBA1* using the 55-bp sequence (Fig. S3B). Because the CRISPR-Cas9 targeting site is more than 50-bp away from the desired editing site, homology dependent repair (HDR) donor vectors with selection markers were used to assist with the enrichment of successfully edited iPSC clones. To correct both *GBA1* alleles, we built two HDR donor vectors: one containing GFP and a puromycin resistance gene as the dual selection markers, while the other had mCherry and a puromycin resistance gene^41–43^. Additionally, the selection markers were flanked with the PiggyBac (PB)-specific inverted terminal repeat sequences (ITRs) and TTAA sequences, facilitating the scarless removal of the selection markers by the excision-only PB transposase, leaving behind the intended correction and only a silent G-to-T mutation in *GBA1* exon 10^41–43^. After the delivery of CRISPR-Cas9, gRNA, and the two HDR donor vectors followed by puromycin selection, ∼3% of iPSCs expressed both GFP and mCherry (P1: GFP+/mCh+) indicating the correction of both *GBA1* alleles (Fig. 3D). Following the transfection of the excision-only PB transposase, ∼2% of GFP+/mCh+ iPSCs lost both GFP and mCherry fluorescence (P2: GFP-/mCh-) reflecting the successful removal of the two selection makers. Sequencing *GBA1* exon 10 using targeted amplicon sequencing revealed that the A-to-G variants in both N370S alleles were indeed reverted to wild-type (WT, *GBA1*: WT/WT) in 4 of the 5 iPSC clones analyzed (Fig. 3E). One WT iPSC clone was chosen as the HT809-WT line and used in subsequent experiments.

To verify the specificity of our *GBA1* editing process, we sequenced the HT809-KO and -WT genome at >30x coverage. When restricting the analysis to variants with a gnomAD allele frequency (AF) of <0.001 and a combined annotation-dependent depletion (CADD) phred score of >30, we detected no unintended variants that were introduced into the HT809-KO or -WT lines (Fig. S3A, KO and WT), demonstrating high specificity of our *GBA1* targeting approach. Because our WGS was based on Illumina short-read sequencing and had reduced read depth and mapping quality at the *GBA1* and *GBAP1* locu*s*^24^, we further amplified full-length *GBA1* and *GBAP1* from all three isogenic lines and sequenced both using PacBio HiFi long-read amplicon sequencing. When demultiplexed long reads were aligned to *GBA1* and *GBAP1*, it was clear that no unintended modifications were introduced into *GBA1* or *GBAP1* in either of the HT809-KO and -WT lines (Fig. 3F and S3C).

On day 35 of DAN differentiation, the HT809-N370S, -KO, and -WT lines all demonstrated similar DAN differentiation efficiency, with the percentage of tdTomato+ cells ranging from ∼70 to 80% as quantified by FACS. This consistency facilitated further direct comparisons among DANs differentiated from the three lines (Fig. S4A).

### Characterization of GCase in the HT809 isogenic lines

Western blot analysis of HT809 isogenic iPSCs and DANs revealed a total loss of GCase in KO iPSCs and DANs (Fig. 4A). GCase levels were reduced in N370S/N370S DANs compared to WT DANs, but interestingly this difference was not seen in corresponding iPSCs (Fig. 4A). Glycosylation analysis with Endoglycosidase H (Endo H) or PNGase F treatment of cell lysates demonstrated that ∼30% of N370S GCase was retained in the ER in iPSCs, and the ER-retained fraction was dramatically reduced in DANs (Fig. 4B). *In vitro* GCase activity measurement using 4-Methylumbelliferyl β-D-glucopyranoside (4-MU-β-Glc) revealed higher GCase activity in DANs than in iPSCs, which was attributable to the higher GCase levels in DANs. HT809-N370S had reduced GCase activity in both DANs and iPSCs compared to HT809-WT, whereas HT809-KO had no activity (Fig. 4C). GCase immunostaining demonstrated the lysosomal localization of both WT and N370S GCase in DANs, but N370S/N370S DANs had reduced levels of GCase (Fig. 4D). GCase staining was barely detectable in KO DANs, confirming the specificity of hGCase-1/23 antibody^44^ (Fig. 4D). In contrast, the expression levels of LIMP2 and Saposin C were similar in all three *GBA1* isogenic DANs (Fig. 4E). Lysosomal GCase activity in live DANs measured using LysoFQ-GBA^45^ further confirmed reduced lysosomal GCase activity in N370S/N370S DANs, with none detected in KO DANs (Fig. 4F).

**Figure 4:**
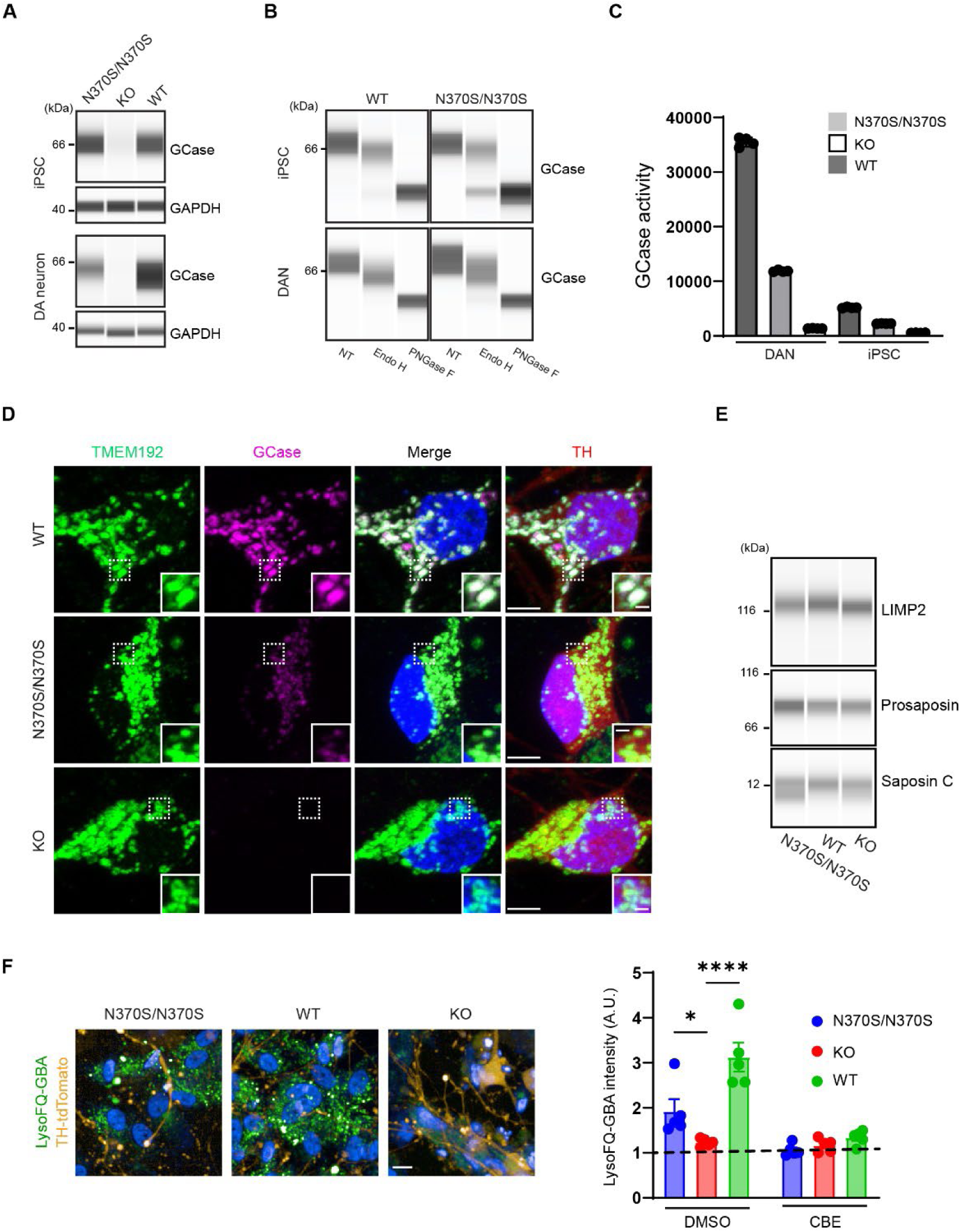
GCase characterization in *GBA1* isogenic lines. **A**. GCase protein levels in isogenic iPSCs and DANs. **B.** GCase glycosylation analysis with Endo H and PNGase F digestion in WT and N370S/N370S iPSCs and DANs. **C.** GCase activity levels in isogenic iPSCs and DANs. **D.** GCase localization in isogenic DANs. Scale bar, 5 µm; insert scale bar, 1 µm. **E.** Levels of LIMP2, prosaposin, and saposin C in isogenic DANs. **F.** Lysosomal GCase activity in isogenic DANs measured using LysoFQ-GBA. Scale bar, 10 µm. Intensities measured in the presence of GCase inhibitor CBE represent the fluorescence of TMEM192-GFP-3xHA in isogenic DANs.

### Glucosylceramide and glucosylsphingosine levels in HT809 isogenic lines

Quantification of GlcCer and GlcSph by supercritical fluid chromatography (SFC) coupled with tandem mass spectrometry (MS/MS) revealed that in WT DANs, the most abundant GlcCer species were C16-, C18-, C20-, C22-, C22:1-, C24-, and C24:1-GlcCer, with levels greater than 0.1 pmole/nmole Pi. In contrast, C14-, C18:1-, C20:1-, C26-, and C26:1-GlcCer, as well as GlcSph, were present in much lower concentrations (<0.1 pmole/nmole Pi). In KO DANs, GlcCer levels for all acyl chain lengths (except C14) increased approximately 6 to 20-fold, with C14-GlcCer remaining below the detection limit. Although GlcSph was only detected at around 0.006 pmole/nmole Pi in WT DANs, its levels in KO DANs surged ∼20,000-fold, reaching 179.23 pmole/nmole Pi. In N370S/N370S DANs, GlcCer levels for all acyl chain lengths (except C14-) increased by about 10 to 50%, while GlcSph increased ∼17.5-fold to 17.5 pmole/nmole Pi (Fig. 5). Compared to DANs, in iPSCs the impact of the N370S GCase and the KO on GlcCer and GlcSph levels was less pronounced. Notably, GlcSph levels in N370S/N370S and WT iPSCs were similar, and only a ∼40-fold increase was observed in KO iPSCs (Fig. S4).

**Figure 5:**
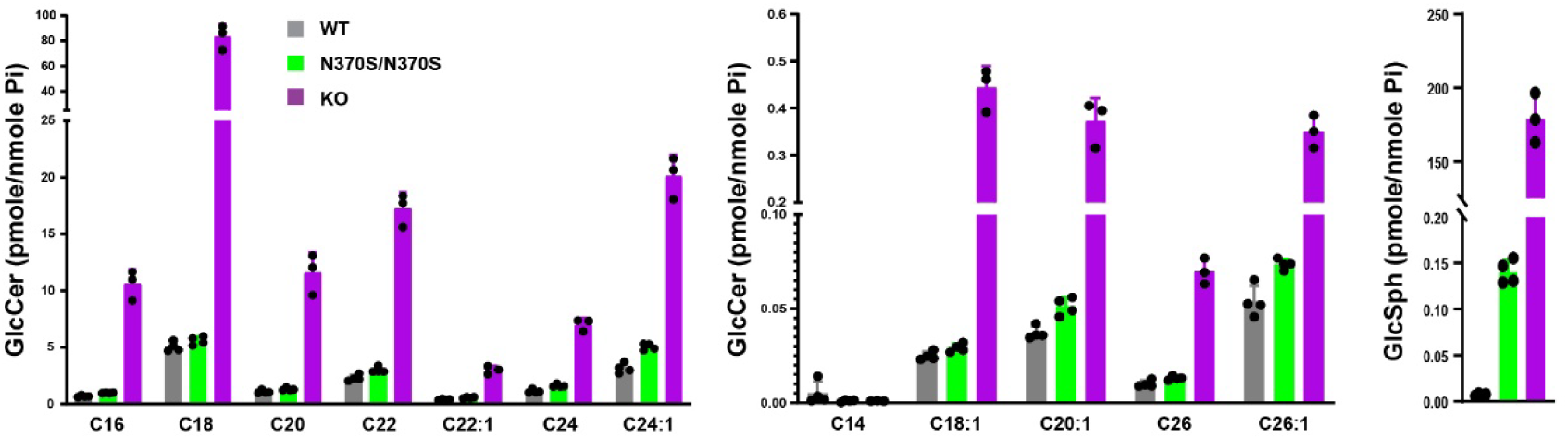
Levels of GlcCer and GlcSph in HT809 isogenic DANs. GlcCer and GlcSph quantification by SFC-MS/MS in *GBA1* isogenic DANs. Phosphate (Pi) amounts were determined in all samples for data normalization.

### Distinct GPNMB phenotypes in N370S/N370S and KO DANs

To specifically probe lysosomal changes in DANs, lysosomes were purified on day 65 of differentiation using LysoIP, and protein abundance was quantified using label-free proteomics. When compared to WT lysosomes, GCase was the most substantially reduced protein in KO lysosomes, followed by cathepsin F (CTSF) and glycoprotein nonmetastatic melanoma protein B (GPNMB) (Fig. 6A). In N370S/N370S lysosomes, GCase levels were reduced by ∼50% and CTSF was also reduced. However, N370S/N370S lysosomes showed a small but significant increase in GPNMB levels (Fig. 6B).

**Figure 6:**
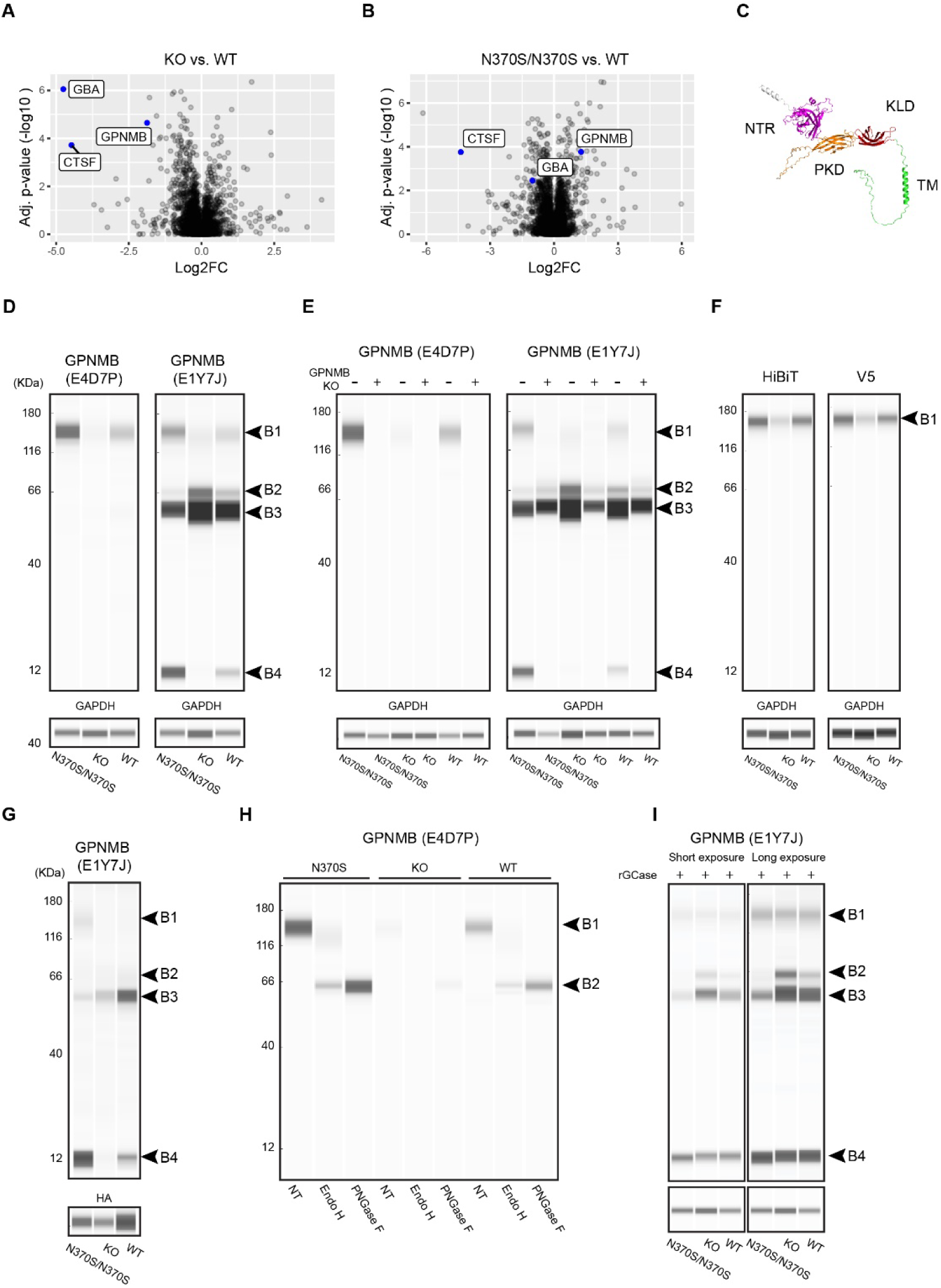
Distinctive GPNMB phenotypes in N370S/N370S and KO DANs. A,B. Volcano plots comparing KO and WT lysosomes (**A)**, and N370S/N370S and WT lysosomes **(B**). **C**. GPNMB structure predicted by ALPHAFOLD. **D.** GPNMB levels in DANs. **E.** Characterization of GPNMB antibody specificity using GPNMB KO DANs. **F.** Expression of HiBiT-GPNMB-V5 in DANs. **G.** Enrichment of GPNMB fragment B4 in lysosomes. **H.** GPNMB glycosylation analysis with E4D7P. **I.** GPNMB expression in isogenic DANs after treatment with recombinant GCase.

**Figure 7:**
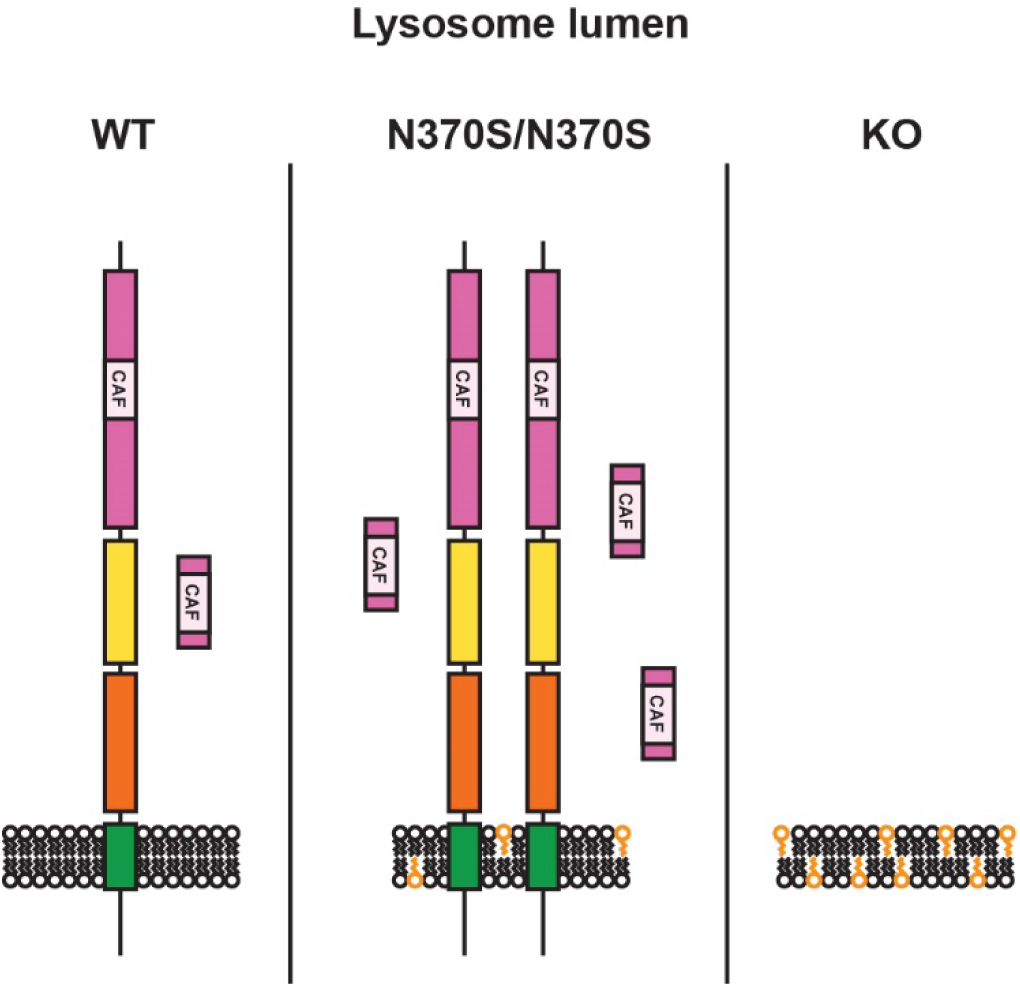
Working model illustrating the levels of GPNMB in *GBA1* isogenic DANs. In WT DANs a portion of GPNMB is processed to a 12-kDa fragment that is enriched in lysosomes. Levels of GPNMB, including both the full-length protein and the 12-kDa fragment, are elevated in N370S/N370S DANs but reduced in KO DANs.

GPNMB is a type 1 transmembrane glycoprotein with several domains: the extracellular domain contains a N-terminal region (NTR) with a predicted core amyloid fragment (CAF)^46^, followed by a polycystic kidney disease (PKD) domain and a Kringle-like domain (KLD); the intracellular domain includes a lysosomal sorting motif (Fig. 6C, S5A). Three unique GPNMB peptides were detected in DAN lysosomes, mapping to the NTR and PKD domains (MS peptide 1, 2, and 3; Fig. S5A). Western blotting analysis of DAN lysates using the monoclonal antibody E4D7P, which recognizes the N-terminal region of GPNMB, identified a single band (∼120 kDa, B1), which is consistent with the molecular weight reported for glycosylated GPNMB (Fig. 6D)^28^. GPNMB levels determined using the E4D7P antibody were lower in KO and higher in N370S/N370S compared to WT DANs. Another monoclonal antibody, E1Y7, which recognizes GPNMB residues around Asp247 (potentially part of the CAF; Fig. S5A), detected B1 as well as several additional bands (∼66 kDa, B2 and B3; ∼12 kDa, B4) (Fig. 6D). As observed with E4D7P, B1 detected by E1Y7 was reduced in KO DANs and increased in N370S/N370S DANs. The intensity of B4 followed a similar trend, being barely detectable in KO DANs. In contrast, B2 showed the opposite trend, with its intensity decreased in N370S/N370S DANs and increased in KO DANs.

To evaluate the specificity of the additional bands detected by E1Y7 (i.e. B2, B3 and B4), GPNMB was knocked out in all three *GBA1* isogenic iPSC lines. Comparison of GPNMB WT and KO DANs lysates using both E4D7P and E1YJ showed that GPNMB KO eliminated B1 and B4, confirming they were derived from GPNMB (Fig. 6E). GPNMB KO also reduced the intensity of B2, particularly in KO and WT DANs, suggesting it originated primarily from GPNMB, though some background remained in the GPNMB KO DANs. B3 remained largely unchanged in GPNMB KO DANs, indicating it was likely non-specific (Fig. 6E). When HiBiT-GPNMB-V5 was overexpressed via lentivirus transduction, the intensity of B1 and B2 increased, further supporting their specificity (Fig. S5C). In contrast, B4 was reduced in HiBiT-GPNMB-V5 transduced DANs, suggesting the cleavage process leading to its production may be sensitive to GPNMB expression levels (Fig. S5C). B3 remained mostly unchanged, further supporting that it is non-specific (Fig. S5C). When HiBiT-GPNMB-V5 transduced DAN lysates were analyzed with HiBiT and V5 antibodies, neither antibody detected B4, suggesting that this fragment lacks both termini (Fig. 6F). Additionally, when LysoIP lysosomes were analyzed using E1Y7J, it was found that B4 was enriched in lysosomes (Fig. 6G). Two neighboring peptides, MS peptide 2 and 3, were detected when the protein gel slice between 10 and 12 kDa was analyzed with Mass Spectrometer (Fig. S5C). Notably, MS peptide 3 overlaps with the antigen recognized by E1Y7J, suggesting that B4 represents a small fragment spanning the overlap region (Fig. S5A). Furthermore, glycosylation analysis using E4D7P, E1Y7J, HiBiT, and V5 antibodies all determined that B1 represented glycosylated GPNMB, whereas B2 represented non-glycosylated GPNMB (Fig. 6H, S5D, and S5E). This data indicated that GPNMB glycosylation was impeded in KO DANs. The distinct pattern of GPNMB expression and processing in N370S/N370S, KO, and WT DANs was less prominent once DANs were treated with recombinant GCase to reduce lysosomal lipid accumulation (Fig. 6I), indicating that the GPNMB phenotypes in N370S/N370S and KO DANs were primarily driven by lysosomal lipid accumulation.

## Discussion

Variants in *GBA1* are the most common genetic risk factor for PD and DLB, and in affected individuals they appear to modify the age of onset and disease progression^13,15^. α-Synuclein accumulates in Lewy bodies (LB) and Lewy neurites (LN) in these disorders, and its accumulation is believed to play a role in disease pathogenesis^47^. Both gain- and loss-of-function hypotheses have been proposed to explain the relationship between *GBA1* variants and α-synuclein accumulation^48,49^, although both theories have shortcomings.

To investigate the PD risk attributable to mutated GCase and to accumulated lipid substrate, we have generated a series of isogenic iPSCs. Unlike previous studies where isogenic lines were generated either by introducing variants into an iPSC line derived from a healthy individual^50^, or by correcting *GBA1* variants in heterozygote lines, we elected to begin from a Parkinson patient iPSC line homozygous for the mild N370S *GBA1* variant to allow for bidirectional modulation of GCase levels and lipid accumulation in a “GD/PD-permissive” genetic background. Using this parental line we then created a *GBA1* KO line to eliminate misfolded GCase and intensify lipid accumulation, and a WT line by reverting the variants on each allele. Because *GBA1* and its highly homologous pseudogene, *GBAP1*, both generate protein-coding transcripts^24^, it was essential to specifically edit *GBA1* in isogenic iPSC lines (i.e. avoiding unintended changes to *GBAP1*) so that the function of *GBA1* could be delineated without interference from *GBAP1*. After evaluating multiple gRNAs targeting *GBA1* exon 4, we identified one that specifically introduced frame-shift mutations in *GBA1*, leading to the generation of the KO line. The identical sequences of *GBA1* and *GBAP1* flanking the *GBA1* N370S point mutation posed a challenge for precise mutation reversion. To overcome this, we targeted CRISPR-Cas9 to a nearby 55-bp genomic sequence unique to *GBA1*. We also used large donor vectors with selection markers to enrich for correctly edited cells. Through both short-read and PacBio HiFi long-read amplicon sequencing, we confirmed successful *GBA1* knockout and the correction of the homozygous N370S mutation to WT without altering *GBAP1*. Whole-genome sequencing (WGS) further confirmed the absence of off-target events. The dramatic accumulation of GlcCer and GlcSph in KO DANs suggests that lysosomal GCase, encoded by *GBA1*, plays the crucial role in catabolism of these lipids. Proteins encoded by *GBA1P*, therefore, most likely function in other pathways.

N370S GCase exhibited reduced stability specifically in DANs but not in iPSCs and showed more ER retention in iPSCs. This suggests that in DANs, N370S GCase may undergo more efficient degradation via ER-associated degradation (ERAD), while some of it successfully folds and traffics to lysosomes. Although patients homozygous for N370S present with non-neuronopathic type 1 GD and only rarely go on to develop PD at advanced ages, our data demonstrate that N370S/N370S DANs accumulated GCase substrates, particularly GlcSph, which exhibited levels 17.5 times higher than WT, whereas GlcCer levels were only ∼10-50% higher. These findings are consistent with previous studies showing a more than 200-fold increase in GlcSph in plasma samples from some patients with type 1 GD, while GlcCer increased by only a factor of 3^51^. This highlights that GlcSph, but not GlcCer, is a useful biomarker for diagnosing GD and monitoring therapeutic effects.

As the culprit in the loss-of-function hypothesis, GlcCer and GlcSph are demonstrated to cause α-synuclein aggregation by stabilizing soluble oligomeric intermediates^21–23^ and accelerating its aggregation into oligomeric and/or fibrillary states^52^. Here, lysosomal proteomics analyses revealed that these lipids regulate GPNMB levels in DANs. The N-terminal region of GPNMB was processed to produce a 12-kDa fragment (B4) that was enriched in lysosomes. Because the 12-kDa fragment is in the middle of the GPNMB luminal domain, it is reasonable to speculate that at least three fragments are derived from the luminal domain, and that some of these fragments could play functional roles in lysosomes, similar to what has been seen with PMEL fragments^31^. GlcCer and GlcSph accumulation in KO DANs reduced the production of the 12-kDa fragment, probably by impairing GPNMB glycosylation or its trafficking from the ER to the Golgi, as indicated by reduced levels of glycosylated GPNMB (B1). Remarkably, the levels of GPNMB, including glycosylated GPNMB and the 12-kDa fragment, were increased in N370S/N370S DANs, counter to the trend in KO DANs. Importantly, these GPNMB phenotypes in isogenic DANs were diminished after treatment with recombinant GCase to decrease lipid accumulation. This rescue reveals that the GPNMB phenotypes in N370S/N370S and KO DANs were driven primarily by the different levels of lipid accumulation.

*GPNMB* is considered a PD risk gene where the PD risk–associated haplotype leads to three-fold higher *GPNMB* expression^25,26,53^ and elevated GPNMB levels in CSF^54^. The specific association of PD risk levels of GPNMB with the mild lipid accumulation seen in DANs homozygous for N370S, a mild *GBA1* variant, potentially establishes a critical molecular pathway in DANs in *GBA1*-PD etiology. This pathway has the potential to help explain PD risk associated with other mild *GBA1* variants, such as E326K (p.E365K), which is prevalent in PD cases^13^ yet does not cause GD^55^, and the recently identified African ancestry-specific *GBA1* noncoding variant that interferes with the splicing of functional *GBA1* transcripts, resulting in slightly reduced GCase levels and activity^14^. The full function of GPNMB and the molecular mechanisms by which its increased levels cause PD risk, are under active research. Neuronal GPNMB was shown to act as a potential receptor for α-synuclein, facilitating its uptake and aggregation in neurons^26^. However, *Gpnmb* KO failed to prevent the spread of alpha-synuclein or loss of dopaminergic neurons in several murine PD models^56^, suggesting that the role of neuronal GPNMB in PD pathogenesis may be more complex. In macrophages, *Gpnmb* was identified as genetic modifier of lysosome function, modulating autophagic flux and cholesterol ester clearance^57^; Gpnmb also recruits LC3 to phagosomes and facilitates lysosome fusion with phagosomes^58^; and Gpnmb loss-of-function increases lysosome pH^59^. Interestingly, a variety of lysosome stressors, such as the lysosomal damaging agent LLOME, the lysosomal V-ATPase inhibitor Bafilomycin A1, and the lysosomal neutralizing agent chloroquine, induce GPNMB expressions and its redistribution to lysosomes and subsequent secretion^58,60,61^. Additionally, in a genome wide association study (GWAS) to identify genetic factors that modify the life span of 15 inbred mouse strains injected with the specific GCase inhibitor conduritol β-epoxide (CBE) to model GD, the DBA/2J strain, lacking Gpnmb due to a spontaneous mutation *Gpnmb*^R150X^,^57^ was one of the strains with the shortest survival, suggesting Gpnmb may impact GD severity^62^.

While GPNMB is observed in DANs containing neuromelanin the substantia nigra^56^, in PD its expression is induced particularly in midbrain microglia^60^. This is also seen in several models of DAN toxicity, including the 1-methyl-4-phenyl-1,2,3,6-tetrahydropyridine (MPTP) and 6-hydroxydopamine (6-OHDA) models, as well as in AAV- and pre-formed fibril (PFF)-induced models of α-synucleinopathies^56^. Microglial GPNMB induction is a feature shared by the PS2APP/TauP301L mouse model of Alzheimer disease^56^, the brain of *Grn* knockout mice, a model for both frontotemporal dementia (FTD) and neuronal ceroid lipofuscinosis-11 (CLN11)^63^, and mouse models of epilepsy^64^. Furthermore, it is seen in the brains of patients with AD^56^ and FTD-*GRN*^63^ suggesting that it is a trait associated with microglia activated by neurodegeneration.

In patients with GD, GPNMB is established as a biomarker for glycosphingolipid (GSL) accumulation. GPNMB levels in cerebrospinal fluid (CSF) correlate with the severity of neuronopathic forms of GD, indicating its cleavage and secretion into the CSF in the brain^65^. In fact, the two peptides identified in CSF match MS peptides 2 and 3 reported in this study. The increased levels of GPNMB in brain and CSF are also evident in *Gba*^flox/flox^; nestin-Cre mice, a model in which GCase deficiency is restricted to neurons and microglia, providing further support that GPNMB in the CSF originates directly from the brain^65^. Additionally, GPNMB levels are elevated in GD patient spleen, particularly in Gaucher cells (i.e. lipid-laden macrophages), and in GD patient plasma, where its levels correlate well with plasma GlcSph levels. These levels are reduced after treatment with enzyme replacement therapy (ERT)^66^. In this study, we found that GPNMB is also a biomarker for GSL accumulation in DANs. A fundamental question in *GBA1*-PD is whether one *GBA1* variant compromises GCase function adequately to cause lysosomal storage of GSL, as in GD^67–70^. Our data suggests that GPNMB can serve as a biomarker for such storage in DANs. Importantly, the cleavage pattern in DANs documented here has major implications for using secreted GPNMB in biofluids such as plasma and CSF to reflect GSL storage in DANs. Since GPNMB induction occurs in activated microglia in PD, microglia are likely the main source of CSF GPNMB. However, the cleavage pattern of GPNMB in these cells has yet to be established. Two recent studies measured GPNMB levels in plasma^71^ and CSF^54,71^ from patients with Lewy body disorders (LBD), with or without *GBA1* variants, but neither found significant difference in GPNMB levels in patients with LBD with and without *GBA1* variants. This might suggest that one *GBA1* variant alone does not lead to lysosomal GLS accumulation in DANs or microglia. However, this interpretation comes with caveats, as neither the commercial GPNMB ELISA kit^71^ nor the GPNMB SOMAmer^54^ used to measure plasma and CSF GPNMB has been characterized for their detection of GPNMB fragments produced by DANs or microglia.

In summary, we discovered the bidirectional regulation of GPNMB by GCase deficiency in *GBA1* isogenic DANs. Associating PD-casual causal levels of GPNMB with lipid levels specific to N370S, a mild GBA1 variant, potentially establishes a critical molecular pathway in GBA1-PD etiology, helping explain PD risk associated with other mild GBA1 variants. While the current study focused on GPNMB phenotypes, there were other relevant leads identified that merit further evaluation. Moreover, this unique panel of *GBA1* isogenic iPSC lines, engineered from a GD/PD background, will facilitate drug discovery efforts targeting *GBA1* variants and enable genetic screens to further identify and characterize factors modulating GCase activity.

## Author Contributions

C.C. and R.S. conducted genetic engineering of the iPSC lines. C.C., C.M., and E.H. conducted most of the experiments including iPSC differentiation, GCase levels and activity, LysoIP, Immunofluorescence, etc. G.P. genotyped *GBA1* in the patient. Y.H., Z.L., I.K., K.A., and Y.Q conducted cellular proteomics and analyzed the data. Y.L. performed mass spectrum analysis of purified lysosomes and protein gel slice. X.J. performed lipidomics analysis and analyzed the data. D.W. and M.H. conducted high-content imaging and analyzed the data. M. P. conducted PacBio HiFi long-read sequencing and analyzed the data. J.L. and C.B. conducted WGS variant and PRS analysis. E.S. and Y.C. supervised the project and wrote the manuscript.

## Data and Materials Availability

Raw fastq files and proteomics dataset will be available in the ASAP CRN. GBA1 isogenic iPSC lines will be available upon request.

## Methods

### Molecular biology

The CLYBL-targeting pC13N-iCAG.copGFP vector (Addgene, 66578) was used as a backbone to generate pC13N-TMEM192-mGFP-3xHA-T2A-TOM20-mTagBFP-TwinStrep to enable the expression of the transgenes under the control of a constitutive chimeric CAG promoter^39^. A custom gene block (GENEWIZ) encoding TMEM192-eGFP-3xHA_T2A_Tom20-BFP-twinStrep replaced the copGFP sequence in the original plasmid through *SpeI* and *MluI* digestion and Hi-T4™ ligation (New England Biolabs, M2622). The HDR donor vectors used to revert *GBA1* N370S to WT were built using pDONOR-tagBFP-PSM-EGFP (Addgene, 100603) and pDONOR-tagBFP-PSM-dTOMATO (Addgene, 100604) as backbones. After the two vectors were digested with *Hpa*I, two custom gene blocks corresponding to the left 1-kb HDR arm and the right 1-kb HDR arm were assembled into each via NEBuilder® HiFi DNA Assembly (New England Biolabs, E5520), thus generating the *GBA1-*WT-pDONOR-tagBFP-PSM-EGFP and *GBA1-*WT-pDONOR-tagBFP-PSM-dTOMATO. Excision-only PiggyBac Transposase expression vector (PB220PA-1) was purchased from System Biosciences.

### Collecting and genotyping patient fibroblasts and reprogramming into iPSCs

The patient HT809 was evaluated under the IRB-approved clinical protocol 86HG0096 at the National Institutes of Health and provided informed consent prior to participation. Sanger sequencing of *GBA1* was performed as previously described^11^. Fibroblasts were obtained by punch biopsy and cultured in Dulbecco’s Modified Eagle’s Medium (DMEM) supplemented with 15% fetal bovine serum (FBS; R&D systems, G22133) and with 1× penicillin-streptomycin (ThermoFisher Scientific, 15140-122) at 37 °C. The CytoTune™-iPSC 2.0 Reprogramming System (ThermoFisher Scientific, A16517), based on non-integrative Sendai virus, was used to generate iPSCs at the National Heart, Lung and Blood Institute’s iPSC Core according to standard procedures.

### iPSCs maintenance and DAN differentiation

iPSCs were maintained in Essential 8 Medium (ThermoFisher Scientific, A1517001) on Vitronectin-coated plates (ThermoFisher Scientific, A14700;) without the addition of antibiotics. Media was exchanged daily, and cells were kept at 37 °C and 5% CO^2^. Cells were passaged by dissociation in 0.5 mM EDTA every 3-4 days and resuspended in Essential 8 Medium supplemented with 10 μM Y-27632 dihydrochloride (R&D Systems, 1254). The DAN differentiation protocol was adopted from Kim et.al. with minor adaptations^36^. iPSCs were dissociated into single cells with Accutase and plated on Geltrex-coated plates (ThermoFisher Scientific, A1413302) at 400,000 cells/cm^2^ in Neurobasal Medium (ThermoFisher Scientific, 21103-049) with 1× GlutaMAX (ThermoFisher Scientific, 35050079), 1× N2 (ThermoFisher Scientific, 17502048), and 1× B-27 without Vitamin A (ThermoFisher Scientific, 12587010,). The media was supplemented with 250 nM LDN 193189 dihydrochloride (R&D Systems, 6053), 2 μM A 83-01 (R&D Systems, 2939), 200 ng/mL human Sonic Hedgehog/Shh (C24II) N-Terminus (R&D Systems, 1845-SH-500), 2 μM SAG 21k (R&D Systems, 5282), 0.7μM CHIR99021(R&D Systems, 4423), 10μM Y-27632 (R&D Systems, 1254), and 5 μM Emricasan (R&D Systems, 7310). On day 4, CHIR99021 was increased to 3 μM. On days 8 and 9, cells were cultured in Neurobasal/GlutaMAX/1X N2/B-27 supplemented with 3 μM of CHIR99021. On day 10, media was changed to Neurobasal/GlutaMAX/B-27 with 20 ng/ml brain-derived neurotrophic factor (BDNF, Peprotech, 450-02), 20 ng/ml glial cell line-derived neurotrophic factor (GDNF, Peprotech, 450-10), 200 μM ascorbic acid (Sigma-Aldrich, 4034), 1 ng/ml transforming growth factor type β3 (TGFβ3, R&D Systems, 243-B3), 200 μM dibutyryl cAMP (Sigma-Aldrich, D0627) and 3 μM CHIR99021. On day 11 cells were sub-plated at 800,000 cells/cm^2^ with Accutase in the same media with Y-27632 and Emricasan. On day 13, CHIR99021 was substituted with 10 μM DAPT (R&D Systems, 2634). Cells were dissociated for the last time on day 25 and seeded at 200,000 cells/cm^2^. Subsequently, the media was changed twice each week until the DANs were collected between day 45 and 75.

### iPSC gene editing with TALEN, CRISPR-Cas9, and piggyBac transposase

To insert pC13N-TMEM192-mGFP-3xHA-T2A-TOM20-mTagBFP-TwinStrep into the *CLYBL* safe harbor site, HT809-N370S iPSCs were transfected with pC13N-TMEM192-mGFP-3xHA-T2A-TOM20-mTagBFP-TwinStrep together with two TALEN constructs targeting *CLYBL* (pZT-C13-L1, Addgene 62196, and pZT-C13-R1, Addgene 62197). After GFP+ iPSC clones were derived, the expression of TMEM192-mGFP-3xHA was validated with Western blots.

Gene editing with CRISPR–Cas9 and piggyBac transposase was carried out based on the protocol developed by the Allen Institute for Cell Science with some modifications^72,73^. The NEON Transfection system (ThermoFisher Scientific) was used to deliver into iPSCs the CRISPR-Cas9 machinery including recombinant Cas9 protein and sgRNA, the donor plasmids, and the plasmid expressing excision-only piggyBac transposase. To generate the KO line, iPSCs were dissociated into single cells using TrypLE and a cell pellet of 8 × 10^5^ cells was resuspended in 100 μl Neon Buffer R supplemented with 54 pmol of TrueCut Cas9 protein v2 (ThermoFisher Scientific, A36497,) and 54 pmol of TrueGuide 1-piece modified synthetic guide RNA (5′-cgacggatggagctgagtat-3′). The Cas9–sgRNA ribonucleoprotein (RNP) was precomplexed for a minimum of 10 minutes at room temperature before addition to the cell suspension. Electroporation was done with 1 pulse at 1,300 V for 30 ms. Cells were then immediately plated onto LN521-coated 6-well dishes with StemFlex medium and CEPT^74^ was applied for the initial 24 hours after plating. When transfected cells had recovered to approximately 70% confluence (usually after 3–4 days), cells were collected for FACS using TrypLE. The cell suspension (1.0 × 106 cells/ml in StemFlex with CEPT) was filtered through a 35-μm mesh filter into polystyrene round-bottomed tubes. Single cells were deposited into LN521-coated 96-well plates that were filled with CEPT-containing StemFlex medium. Half-medium changes were performed with StemFlex medium (without CEPT) on day 3, day 6, and day 9 after FACS sorting. Single-cell clones were transferred to 6-well plates for continued culture and further analysis on day 12.

To generate the reverted WT line, iPSCs were first electroporated with 54 pmol of TrueCut Cas9 protein v2, 54 pmol of TrueGuide 1-piece modified synthetic guide RNA (5′-cttgccctgaaccccgaagg-3′), 2 μg donor plasmids (equal amount of the GFP+ HDR donor vector and the mCherry+ one). After puromycin selection, GFP+/mCherry+ iPSCs were enriched by FACS. GFP+/mCherry+ iPSCs were then electroporated with a plasmid expressing excision-only piggyBac transposases. After recovery for 5 days, iPSCs were collected for FACS and GFP-/mCherry-iPSCs were deposited into 96-well plates for single-cell cloning. Established clones were further analyzed to identify those carrying the correct reversion.

### Karyotyping

iPSC clones were dissociated and plated at 1×10^6^ cells/well in 6 well plates in StemFlex with 10 μM Y-27632. Colcemid was added to the cells (1:100, Roche, 10295892001) 24 hours later, and cells were karyotyped using standard procedures at the National Human Genome Research Institute Cytogenetics Core Facility.

### Whole genome sequencing

PCR-free libraries were prepared from 1 microgram genomic DNA using the TruSeq® DNA PCR-Free HT Sample Preparation Kit (Illumina). The median insert size was approximately 400 bp. Libraries were tagged with unique dual index DNA barcodes to allow pooling of libraries and to minimize the impact of barcode hopping. Libraries were pooled for sequencing on the NovaSeq 6000 (Illumina) obtaining at least 300 million 151-base read pairs per individual library. NVIDIA Parabricks GATK human_par pipeline was used for variant calling. Paired-end reads that passed standard Illumina quality filters for a sample, were aligned in fastq format to the human reference genome sequence hg19 (GRCH 37) using the bwa-mem v.0.7.15 component of the NVIDIA Parabricks pipeline. The aligned bam file was sorted, and the duplicates marked using MarkDuplicates. After Base Quality Score Recalibration, variant calling was performed by the Haplotype Caller - v4.1.0.0 generating a GVCF with both reference and variant calls.

Polygenic risk scores for AD and PD were calculated using PLINK (v1.9) with the weights of recent GWAS. As a reference population for the polygenic risk score, we used AD^75^ (Data-Field 131,037), PD^76^ (Data-Field 131,023) and control (no known neurodegenerative disease, no parent with a known neurodegenerative disease and ≥60 years old at recruitment) samples from the UK Biobank (application ID: 33,601).

### Short-read amplicon sequencing

Two pairs of primers were designed to generate amplicons from *GBA1* exon 4 (amplicon size: 416 bp; EZ-Exon4-Forward: cagatgtgtccattctccatgtcttc and EZ-Exon4-Reveres: ctggaacttctgttctggctgc) and exon 10 (amplicon size: 457 bp; EZ-Exon10-Forward: aactactgaggcacttgcagc and EZ-Exon10-Reverse: tcgtggtgtagagtgatgtaagccand), respectively. Purified amplicons were sequenced at Genewiz using the Amplicon-EZ service to generate ∼50K reads for each sample. Reads were analyzed using Crispresso2 (http://crispresso2.pinellolab.org/)^77^.

### Targeted PacBio HiFi long-read amplicon sequencing

A PacBio library was prepared from 1 µg of amplicon product using the Adapter-barcoded workflow of ‘Preparing multiplexed amplicon libraries using SMRTbell® prep kit 3.0’ (Pacific Biosciences). The library was run on one 8M SMRTCell using Sequel II Sequencing Kit 2.0 sequencing reagents. Sequencing was performed on a Sequel IIe sequencer (Pacific Biosciences) running instrument control software version 11.0.0.144466 with a 30-hour movie collection time and 2-hour pre-extension. Circular consensus sequence (CCS) reads were generated from the initial subread data using the ccs program (version 6.3.0 within PacBio SMRTLink version 11.0.0.146107). The CCS reads were demultiplexed using lima (version 2.5.1/SL v11.0.0.146107) to generate full-length (FL) CCS reads for each sample with the proper 5prime and 3prime barcodes and to remove the barcode primer sequences. Full-length non-chimeric (FLNC) reads were generated from the FL reads using isoseq refine (version 3.5.0/SL v11.0.0.146107) which identifies FL reads with the correct 5prime and 3prime barcode orientation and filters out concatemerized products.

Bamsieve (version 0.2.0 within SMRTLink version 25.1.0.257715) was used to randomly select 2000 FLNC reads from each sample; the downsampled fastq was indexed with samtools (version 1.14) fqidx. Downsampled reads were aligned to GRCh38/hg38 using pbmm2 (version 1.16.0/SL v25.1.0.257715). Consensus sequences were generated from the downsampled reads using pbaa (https://github.com/PacificBiosciences/pbAA, version 1.1.0/SL v25.1.0.257715) cluster. The downsampled reads were aligned against the amplicon target region using pbmm2 and color tags added to the read alignments using pbaa bampaint with the read information from the pbaa cluster step.

### Simple Western Automated Western Blot JESS System

DAN and iPSC pellets were resuspended in the Triton-X buffer (1% Triton X-100, 10% glycerol, 150 mM NaCl, 25 mM HEPES pH 7.4, 1 mM EDTA, 1.5 mM MgCl2, supplemented with one EDTA-free protease inhibitor tablet per 10 mL of buffer (Pierce, A32965), lysed through grinding and 3 cycles of freeze-thaw. After incubation on ice for 30 min, lysates were centrifuged at 21,000 × g for 15 minutes at 4°C. The supernatant was collected as the Triton soluble fraction. The pellets were then solubilized in the SDS buffer (2% SDS, 10 mM Tris, pH 7.5). Protein concentration was determined by BCA (ThermoFisher Scientific, 23225) according to manufacturer’s instructions and absorbance values were detected on FlexStation® 3 Multi-Mode Microplate Reader (Molecular Devices).

Comprehensive instructions for Jess (Bio-Techne, 004-650) are provided by the manufacturer. Runs were set up according to instructions in the EZ Standard Pack 1 (Bio-Techne, PS-ST01EZ-8). In brief, the DTT was prepared with 40 μl of deionized water and the biotinylated ladder with 20 μl of deionized water respectively. Fluorescent 5X Master Mix was prepared with 20 μl of Sample Buffer and 20 μl of the DTT solution. Samples were prepared by combining one-part fluorescent master mix with four parts sample to a final concentration of 1 μg/μl. Samples and a biotinylated ladder were incubated for 5 min at 95 °C. Fluorescence Separation Capillary Cartridges (Bio-Techne, SM-FL001) were loaded according to instructions in the RePlex pack (Bio-Techne, RP-001) with 3 μL sample per well. Primary antibodies were diluted in in antibody diluent 2 and normalization was based on total protein or reference gene. Plates were centrifuged for 5 min at 1000 × g before addition of wash buffer, luminol-peroxide mix and RePlex reagent. Visualization was performed with SW Compass (Simple Western).

### Glycosylation analysis (Endo H and PNGase F assay)

Glycosidase sensitivity analysis was performed as previously described^78^. Briefly, iPSC and DAN pellets were lysed in 1% Triton X-100 lysis buffer supplemented with Protease and Phosphatase Inhibitor Mini Tablets (Pierce, A32959). The cells were sonicated for 10 quick pulses using a Sonic Dismembrator Model D100 (Fisher Scientific), left standing for 15-30 min on ice, and went through three cycles of freezing and thawing before centrifuging for 15 min (13,000 x g; 4°C). The supernatant containing lysate was collected and stored at -80°C until digestion.

For each condition, 40 – 50 µg of protein lysate from each sample was denatured in 1X Glycoprotein Denaturing Buffer (New England Biolabs, B1704S) with 20 µL total volume at 100°C for 10 min. The denatured protein lysates were digested with glycosidase enzymes, Endo Hf (New England Biolabs, P0703S;) or PNGase F (New England Biolabs, P0708L). For Endo Hf digestion, 20 µL of denatured protein lysate was combined with 2.5 µL of 10X GlycoBuffer 3 (New England Biolabs, B1720S) and 2.5 µL of Endo Hf in a total reaction volume of 25 µL. For PNGase F digestion, 20 µL of denatured protein lysate was combined with 2.6 µL of 10X GlycoBuffer 2 (New England Biolabs, B3704S), 2.6 µL of 10% NP-40 (New England Biolabs, B2704S), and 1.5 µL of PNGase F in a total reaction volume of 26.7 µL. Undigested control samples were prepared by combining 20 µL of denatured protein lysate with 5 µL of water. The reactions were incubated at 37°C for 1.5 h. After the reactions were completed, samples were washed with 0.1x Sample Buffer (Simple Western) before being analyzed using Jess. To wash the samples, 480 uL 0.1X Sample Buffer was added to the EndoH/PNGase F reactions mixture to bring the final volume to 500 uL. Samples were then transferred to a 10K Amicon Ultra Centrifugal Filter and centrifuged for 20 minutes to reduce the total volume to ∼20 uL.

### GCase activity assay (4-MUG)

GCase enzymatic activity was determined based on cleavage of the fluorogenic substrate 4-methylumbelliferyl-β-D-glucopyranoside (4-MUG). Briefly, freshly-prepared GCase buffer [McIlvaine buffer (0.2 M Na2HPO4 and 0.1 M citric acid titrated to pH 5.4); cOmplete™, Mini, EDTA-free Protease Inhibitor Cocktail (Roche, 11836170001); 0.25% (v/v) Triton X-100 (Sigma-Aldrich, T9284)] was activated with 0.2% (w/v) sodium taurocholate (Sigma-Aldrich, 86339). Protein was extracted from iPSC and DAN pellets in GCase buffer through pipetting, at least two 10-sec pulsed bath sonication steps (S-4000, Misonix), and three freeze-thaw cycles. The lysates were then centrifuged for 15 min (21,000 x g; 4°C), and the supernatant containing extracted protein was collected. Protein concentration was measured BCA Protein Assay Kit (ThermoFisher Scientific, 23225), and samples were diluted to 1 µg/µL, unless specified otherwise. Final protein concentration was verified by a second BCA assay and used for data normalization.

The 4-MUG GCase activity assay was conducted in a Greiner black 384-well plate (Greiner, 781209). Protein samples were added (10 µL/well) and further diluted with GCase buffer (5 µL/well). The plate was sealed, centrifuged, and incubated (37°C) in a shaking plate incubator at 600 rpm for 15 min. After brief centrifugation, 15 µL of assay buffer (2.5 mM 4-MUG; 0.25% v/v DMF in GCase buffer) was added to each assay well. The plate was sealed, centrifuged, and incubated (37°C) for 60 min, shaking at 450 rpm. After incubation, the plate was centrifuged, and 30 µL of 1 M glycine stop solution (pH 10.5) was added to each well. Fluorescence was top read on a FlexStation® 3 Multi-Mode Microplate Reader (Molecular Devices; 365 nm excitation wavelength; 449 nm emission wavelength; 435 nm cutoff; 6 reads/well).

### Lysosomal GCase activity assay using LysoFQ-GBA

Day 25 DANs were seeded in Geltrex-coated 96-well, black, optically clear, flat-bottom, tissue-culture treated PhenoPlates (PerkinElmer) at 5,000 cells/well and lysosomal GCase activity was determined on Day 35 following an established protocol^79^. Briefly, DANs were incubated with 10 µM LysoFQ-GBA for 2 hours followed by image acquisition using a PerkinElmer Opera Phenix Plus confocal high-content screening system. Images were acquired in both LysoFQ-GBA and tdTomato channels, and tdTomato channels were used as masks to measure LysoFQ-GBA intensities in DANs. Baseline intensity in the LysoFQ-GBA channel due to Tmem192-GFP-3xHA expression was determined by inhibiting GCase activity using Conduritol B epoxide (CBE) pretreatment for 1 hour.

### Immunocytochemistry

DANs were seeded in Geltrex-coated 8-well imaging chambers (Ibidi, 80807-90) at differentiation day 25 at a density of 20,000 cells/cm^2^. For immunostaining, cells were washed once with PBS (pH 7.4) and fixed with 4% paraformaldehyde (30 min, RT). Fixed cells were washed with PBS (3 x 5 min) and incubated overnight (4°C) with primary antibody in ICC antibody diluent [PBS containing 0.05% or 0.1% saponin (filtered) and 1% BSA]. After primary antibody incubation, cells were washed with PBS (3 x 5 min, at least) and incubated for 1 h (RT) with secondary antibody in antibody diluent. After secondary antibody incubation, cells were washed with PBS (3 x 5 min, at least) and underwent nuclear staining. For Hoechst staining, Hoechst 33342 solution (ThermoFisher Scientific,,62249,) was diluted in PBS to 5 µg/mL and added during the third wash. For DAPI staining, cells were stored at 4°C in PBS with NucBlue™ Fixed Cell ReadyProbes™ Reagent (DAPI) (ThermoFisher Scientific,,R37606) until imaging.

### LysoIP from DANs

50×10^6^ iPSCs or 35×10^6^ DANs were used for each Lyso-IP purification. After washing with cold DPBS, cells were scraped off the plate in ice cold KPBS buffer (136 mM KCl, 10 mM KH2PO4, pH 7.25 adjusted with KOH) and spun down at 1000xg for 3 min at 4C in a 15-mL conical tube. Cell pellets were resuspended in 950 uL of KPBS + protease/phosphatase inhibitors and 25 uL of suspension was saved as the input. Cells in suspension were homogenized using a motorized homogenizer 3-5 times each for 10 sec. Homogenates were centrifuged at 1000xg for 2 min at 4C to remove nuclei. Supernatants containing the organelles were transferred to new tubes to mix with HA beads in KPBS + inhibitors for 5 min on a rotor at 4C. After washing three times using 1 mL KPBS + Protease/Phosphatase inhibitors, the HA beads were mixed with 100 uL of Lysis Buffer (2% SDS, 50 mM TEAB pH 8.5) and boiled for 10 min at 100c to elute proteins.

### Label-free proteomics

In-solution samples were reduced with 3 mM Tris(2-carboxyethyl)phosphine hydrochloride (TCEP), alkylated with 5 mM N-Ethylmaleimide (NEM), and digested with trypsin on micro S-Trap columns with 1:10 (w/w) at 37 °C for 18hr. Tryptic digests were cleaned with an Oasis HLB 10mg plate (Waters). ∼1µg of each sample was injected into a nano-LC/MS/MS system where an Ultimate 3000 HPLC was coupled to an Orbitrap Fusion Lumos Tribrid Mass Spectrometer (ThermoFisher Scientific) via an EASY-Spray™ ion Sources (ThermoFisher Scientific). Peptides were separated on an ES902 column. Mobile phase A contained 0.1% formic acid in LC-MS grade water, and Mobile phase B 0.1% formic acid in LC-MS grade acetonitrile. Mobile phase B was increased from 3% to 18% over 60 min, then from 18% to 25% over 9min. LC-MS/MS data were acquired in data-dependent mode. The MS1 scans were performed in orbitrap with a resolution of 120K, a mass range of 375-1500 m/z, and an AGC target of 4 x 10^5^. The quadrupole isolation window was 1.6 m/z. The precursor ion intensity threshold to trigger the MS/MS scan was set at 1 x 10^4^. MS2 scans were conducted in the ion trap. Peptides were fragmented using the HCD method, and the collision energy was fixed at 35%. MS1 scan was performed every 3 sec. As many MS2 scans as possible were acquired within the 3-sec cycle. Proteome Discoverer software version 2.4 was used for protein identification and quantitation. Raw data were searched against the Sprot Human database and a house-built database containing the sequence of IgG used in the pull-down experiment. Up to one missed cleavage was allowed for trypsin digestion. Variable modifications included Oxidation (M), NEM (C), Met-loss (Protein N-term), Met-loss+Acetyl (Protein N-term) and Acetyl (Protein N-term). Mass tolerances for MS1 and MS2 scans were set to 5 ppm and 0.6 Da, respectively. Protein abundance values were calculated for all proteins identified by summing the abundance of unique peptides matched to that protein. The normalization was performed against total peptide abundance. Protein ratios were calculated by comparing the protein abundances between two conditions. p-values were calculated with the ANOVA method.

In-gel bands were reduced with 5 mM TCEP, alkylated with 5 mM NEM, and digested with trypsin. Tryptic digests were extracted from the gel and cleaned with an Oasis HLB microelution plate (Waters). ∼0.2 µg of each sample was injected. The method described above was used for data acquisition. In addition to HCD runs, each sample was also acquired using an EThCD fragmentation method. Raw data were processed with Mascot Distiller and searched with Mascot Daemon software (Matrix Science). The sequences of the GPNMB fragments were added into the house-built database. Data were searched against both Sprot Human and house-built databases. The mass tolerances for precursor and fragment were set to 5 ppm and 0.5 Da, respectively. Trypsin was used as enzyme with up to two missed cleavages allowed. NEM on cysteines was set as fixed modification. Variable modifications included Oxidation (M), Met-loss (Protein N-term), Met-loss+Acetyl (Protein N-term) and Acetyl (Protein N-term). The search results were filtered by a false discovery rate of 1% at the protein level. Peptides detected by database search were manually curated.

### Fluorescence microscopy

Confocal images were acquired with a Zeiss LSM 880 confocal microscope and Airyscan images with a Zeiss LSM 980 confocal microscope, both using a 63x oil objective. High-content imaging was conducted on a PerkinElmer Opera Phenix Plus confocal high-content screening system; images were captured in 27 fields per well with a 63x water, NA 1.15 objective. Image analysis was performed using PerkinElmer Columbus software.

### Lipidomic analysis

Lipid extraction and SFC-MS/MS analysis were performed by the Analytical Unit of the Lipidomics Shared Resource at the Medical University of South Carolina (MUSC) Hollings Cancer Center using supercritical fluid chromatography (SFC) separation coupled with tandem mass spectrometry (MS/MS) detection. DANs were pelleted in ice-cold DPBS and rinsed once with 0.15 M NaCl solution (Cat.#46-032-CV; Corning) and stored at -80°C prior to analysis. Extracted lipid samples and synthetic standards were processed on a setup comprising a Waters Acquity SFC/UPC2 Chromatography System and TSQ Quantum Access Max Triple Quadrupole Mass Spectrometers (ThermoFisher Scientific) operating in positive MRM mode and employing a gradient elution. Quantitation required the generation of analyte-specific eight-point calibration curves (analyte/internal standard peak area ratio versus analyte concentration), as described previously^80,81^. Lipid measurements [pmol/total sample] were normalized to levels of inorganic phosphate [nmol] in each sample, as determined by Bligh & Dyer (B&D) re-extraction of an aliquot from the original total sample extract^82^.

### Flow cytometry

DANs and iPSCs were washed with DPBS (Cat.# 14190144, Thermo Fisher) and dissociated with Accutase at 37 °C. Cells were resuspended in Neurobasal media supplemented with Y-27632 and Emricasan and were analyzed for the fluorescent intensity of TH-tdTomato using BD FACSAria™ Fusion Flow Cytometers.

## References

1. Blauwendraat, C., Nalls, M. A. & Singleton, A. B. The genetic architecture of Parkinson’s disease. Lancet Neurol. 19, 170–178 (2020).

2. Singleton, A. B. et al. alpha-Synuclein locus triplication causes Parkinson’s disease. Science (80-.). 302, 841 (2003).

3. Polymeropoulos, M. H. et al. Mutation in the α-synuclein gene identified in families with Parkinson’s disease. Science (80-.). 276, 2045–2047 (1997).

4. Kim, J. J. et al. Multi-ancestry genome-wide association meta-analysis of Parkinson’s disease. Nat. Genet. 56, 27–36 (2024).

5. Chang, D. et al. A meta-analysis of genome-wide association studies identifies 17 new Parkinson’s disease risk loci. Nat. Genet. 49, 1511–1516 (2017).

6. Pan, H. et al. Genome-wide association study using whole-genome sequencing identifies risk loci for Parkinson’s disease in Chinese population. *npj Park*. Dis. 9, (2023).

7. Sidransky, E. et al. Multicenter Analysis of Glucocerebrosidase Mutations in Parkinson’s Disease. N. Engl. J. Med. 361, 1651–1661 (2009).

8. Nalls, M. A. et al. A multicenter study of glucocerebrosidase mutations in dementia with Lewy bodies. JAMA Neurol. 70, 727–735 (2013).

9. Grabowski, G. A., Gaft, S., Horowitz, M. & Kolodny, E. H. Acid β-Glucosidase: Enzymology and Molecular Biology of Gaucher Diseas. Crit. Rev. Biochem. Mol. Biol. 25, 385–414 (1990).

10. Zhao, H. & Grabowski, G. A. Gaucher disease: perspectives on a prototype lysosomal disease. Cell. Mol. Life Sci. C. 59, 694–707 (2002).

11. Tayebi, N. et al. Gaucher disease with parkinsonian manifestations: Does glucocerebrosidase deficiency contribute to a vulnerability to parkinsonism? Mol. Genet. Metab. 79, 104–109 (2003).

12. Várkonyi, J. et al. Gaucher disease associated with Parkinsonism: Four further case reports. Am. J. Med. Genet. 116 A, 348–351 (2003).

13. Duran, R. et al. The glucocerobrosidase E326K variant predisposes to Parkinson’s disease, but does not cause Gaucher’s disease. Mov. Disord. 28, 232–236 (2013).

14. Álvarez Jerez, P., et al. African ancestry neurodegeneration risk variant disrupts an intronic branchpoint in GBA1. Nat. Struct. Mol. Biol. 31, 1955–1963 (2024).

15. Alcalay, R. N. et al. Comparison of parkinson risk in ashkenazi jewish patients with gaucher disease and gba heterozygotes. JAMA Neurol. 71, 752–757 (2014).

16. Blauwendraat, C. et al. Genetic modifiers of risk and age at onset in GBA associated Parkinson’s disease and Lewy body dementia. Brain 143, 234–248 (2020).

17. Maor, G. et al. The contribution of mutant GBA to the development of Parkinson disease in Drosophila. Hum. Mol. Genet. 25, 2712–2727 (2016).

18. Maor, G., Rapaport, D. & Horowitz, M. The effect of mutant GBA1 on accumulation and aggregation of a synuclein. Hum. Mol. Genet. 28, 1768–1781 (2019).

19. Kuo, S. H. et al. Mutant glucocerebrosidase impairs α-synuclein degradation by blockade of chaperone-mediated autophagy. Sci. Adv. 8, (2022).

20. Baden, P. et al. Glucocerebrosidase is imported into mitochondria and preserves complex I integrity and energy metabolism. Nat. Commun. 14, 1–21 (2023).

21. Mazzulli, J. R. et al. Gaucher disease glucocerebrosidase and α-synuclein form a bidirectional pathogenic loop in synucleinopathies. Cell 146, 37–52 (2011).

22. Fredriksen, K. et al. Pathological α-syn aggregation is mediated by glycosphingolipid chain length and the physiological state of α-syn in vivo. Proc. Natl. Acad. Sci. U. S. A. 118, 1–12 (2021).

23. Zunke, F. et al. Reversible Conformational Conversion of α-Synuclein into Toxic Assemblies by Glucosylceramide. Neuron 97, 92–107.e10 (2018).

24. Gustavsson, E. K. et al. The annotation of GBA1 has been concealed by its protein-coding pseudogene GBAP1. Sci. Adv. 10, (2024).

25. Murthy, M. N. et al. Increased brain expression of GPNMB is associated with genome wide significant risk for Parkinson’s disease on chromosome 7p15.3. Neurogenetics 18, 121–133 (2017).

26. Diaz-Ortiz, M. E. et al. GPNMB confers risk for Parkinson’s disease through interaction with α-synuclein. Science (80-.). 377, (2022).

27. Theos, A. C. et al. The PKD domain distinguishes the trafficking and amyloidogenic properties of the pigment cell protein PMEL and its homologue GPNMB. Pigment Cell Melanoma Res. 26, 470– 486 (2013).

28. Hoashi, T. et al. Glycoprotein nonmetastatic melanoma protein b, a melanocytic cell marker, is a melanosome-specific and proteolytically released protein. FASEB J. 24, 1616–1629 (2010).

29. Berson, J. F. et al. Proprotein convertase cleavage liberates a fibrillogenic fragment of a resident glycoprotein to initiate melanosome biogenesis. J. Cell Biol. 161, 521–533 (2003).

30. Kummer, M. P. et al. Formation of Pmel17 amyloid is regulated by juxtamembrane metalloproteinase cleavage, and the resulting C-terminal fragment is a substrate for γ-secretase. J. Biol. Chem. 284, 2296–2306 (2009).

31. Leonhardt, R. M., Vigneron, N., Rahner, C. & Cresswell, P. Proprotein convertases process Pmel17 during secretion. J. Biol. Chem. 286, 9321–9337 (2011).

32. Lopez, G. et al. Clinical course and prognosis in patients with Gaucher disease and parkinsonism. Neurol. Genet. 2, (2016).

33. Pantazis, C. B. et al. Resource A reference human induced pluripotent stem cell line for large-scale collaborative studies ll Resource A reference human induced pluripotent stem cell line for large-scale collaborative studies. 1685–1702 (2022) doi:10.1016/j.stem.2022.11.004.

34. Blauwendraat, C. et al. Polygenic Parkinson’s Disease Genetic Risk Score as Risk Modifier of Parkinsonism in Gaucher Disease. Mov. Disord. 38, 899–903 (2023).

35. Ahfeldt, T. et al. Pathogenic Pathways in Early-Onset Autosomal Recessive Parkinson’s Disease Discovered Using Isogenic Human Dopaminergic Neurons. Stem Cell Reports 14, 75–90 (2020).

36. Kim, T. W. et al. Biphasic Activation of WNT Signaling Facilitates the Derivation of Midbrain Dopamine Neurons from hESCs for Translational Use. Cell Stem Cell 28, 343–355.e5 (2021).

37. Abu-Remaileh, M. et al. Lysosomal metabolomics reveals V-ATPase- and mTOR-dependent regulation of amino acid efflux from lysosomes. Science (80-.). 358, 807–813 (2017).

38. Xiong, J. et al. Rapid affinity purification of intracellular organelles using a twin strep tag. J. Cell Sci. 132, (2019).

39. Cerbini, T. et al. Transcription activator-like effector nuclease (TALEN)-mediated CLYBL targeting enables enhanced transgene expression and one-step generation of dual reporter human induced pluripotent stem cell (iPSC) and neural stem cell (NSC) lines. PLoS One 10, 1–18 (2015).

40. Woo, E. G., Tayebi, N. & Sidransky, E. Next-Generation Sequencing Analysis of GBA1: The Challenge of Detecting Complex Recombinant Alleles. Front. Genet. 12, 1–5 (2021).

41. Jarazo, J., Qing, X. & Schwamborn, J. C. Guidelines for fluorescent guided biallelic HDR targeting selection with PiggyBac system removal for gene editing. Front. Genet. 10, 1–12 (2019).

42. Arias-Fuenzalida, J. et al. FACS-Assisted CRISPR-Cas9 Genome Editing Facilitates Parkinson’s Disease Modeling. Stem Cell Reports 9, 1423–1431 (2017).

43. Hanss, Z., Boussaad, I., Jarazo, J., Schwamborn, J. C. & Krüger, R. Quality Control Strategy for CRISPR-Cas9-Based Gene Editing Complicated by a Pseudogene. Front. Genet. 10, 1–9 (2020).

44. Jong, T., Gehrlein, A., Sidransky, E., Jagasia, R. & Chen, Y. Characterization of Novel Human β-glucocerebrosidase Antibodies for Parkinson’s Disease Research. J. Parkinsons. Dis. 14, 65–78 (2024).

45. Theka, I. et al. Rapid Generation of Functional Dopaminergic Neurons From Human Induced Pluripotent Stem Cells Through a Single-Step Procedure Using Cell Lineage Transcription Factors. Stem Cells Transl. Med. (2013) doi:10.5966/sctm.2012-0133.

46. Chrystal, P. W. et al. Functional Domains and Evolutionary History of the PMEL and GPNMB Family Proteins. Molecules 26, 1–28 (2021).

47. Spillantini, M. G. et al. α-Synuclein in Lewy bodies. Nature 388, 839–840 (1997).

48. Gegg, M. E., Menozzi, E. & Schapira, A. H. V. Glucocerebrosidase-associated Parkinson disease: Pathogenic mechanisms and potential drug treatments. Neurobiol. Dis. 166, 105663 (2022).

49. Chatterjee, D. & Krainc, D. Mechanisms of Glucocerebrosidase Dysfunction in Parkinson’s Disease: Mechanisms of GBA1-PD. J. Mol. Biol. 435, (2023).

50. Ramos, D. M., Skarnes, W. C., Singleton, A. B., Cookson, M. R. & Ward, M. E. Tackling neurodegenerative diseases with genomic engineering: A new stem cell initiative from the NIH. Neuron 109, 1080–1083 (2021).

51. Dekker, N. et al. Elevated plasma glucosylsphingosine in Gaucher disease: Relation to phenotype, storage cell markers, and therapeutic response. Blood 118, 118–127 (2011).

52. Taguchi, Y. V. et al. Glucosylsphingosine promotes α-synuclein pathology in mutant GBA-associated parkinson’s disease. J. Neurosci. 37, 9617–9631 (2017).

53. Kia, D. A. et al. Identification of Candidate Parkinson Disease Genes by Integrating Genome-Wide Association Study, Expression, and Epigenetic Data Sets. JAMA Neurol. 78, 464–472 (2021).

54. Kaiser, S. et al. A proteogenomic view of Parkinson’s disease causality and heterogeneity. *npj Park*. Dis. 9, 1–13 (2023).

55. Horowitz, M., Pasmanik-Chor, M., Ron, I. & Kolodny, E. H. The enigma of the E326K mutation in acid β-glucocerebrosidase. Mol. Genet. Metab. 104, 35–38 (2011).

56. Brendza, R. et al. Genetic ablation of Gpnmb does not alter synuclein-related pathology. Neurobiol. Dis. 159, 105494 (2021).

57. Robinet, P. et al. Quantitative trait locus mapping identifies the Gpnmb gene as a modifier of mouse macrophage lysosome function. Sci. Rep. 11, 1–11 (2021).

58. Li, B. et al. The melanoma-associated transmembrane glycoprotein Gpnmb controls trafficking of cellular debris for degradation and is essential for tissue repair. FASEB J. 24, 4767–4781 (2010).

59. Tejwani, L., et al. Lysosomes cell autonomously regulate myeloid cell states and immune responses. bioRxiv (2024) 10.1101/2024.11.11.623074.

60. Bogacki, E. C., Longmore, G., Lewis, P. A. & Herbst, S. GPNMB is a biomarker for lysosomal dysfunction and is secreted via LRRK2-modulated lysosomal exocytosis. (2025) 10.1101/2025.01.01.630988.

61. Bogacki, E. C., Lewis, P. A. & Herbst, S. Mutations in GPNMB associated with Amyloid cutis dyschromica alter intracellular trafficking and processing of GPNMB. 0–3 (2023).

62. Klein, A. D. et al. Identification of Modifier Genes in a Mouse Model of Gaucher Disease. Cell Rep. 16, 2546–2553 (2016).

63. Huang, M. et al. Network analysis of the progranulin-deficient mouse brain proteome reveals pathogenic mechanisms shared in human frontotemporal dementia caused by GRN mutations. Acta Neuropathol. Commun. 8, 163 (2020).

64. Liu, M. et al. GPNMB and ATP6V1A interact to mediate microglia phagocytosis of multiple types of pathological particles. Cell Rep. 44, 115343 (2025).

65. Zigdon, H. et al. Identification of a biomarker in cerebrospinal fluid for neuronopathic forms of Gaucher disease. PLoS One 10, 1–11 (2015).

66. Kramer, G. et al. Elevation of glycoprotein nonmetastatic melanoma protein B in type 1 Gaucher disease patients and mouse models. FEBS Open Bio 6, 902–913 (2016).

67. Leyns, C. E. G. et al. Glucocerebrosidase activity and lipid levels are related to protein pathologies in Parkinson’s disease. NPJ Park. Dis. 9, 74 (2023).

68. Surface, M. et al. Plasma Glucosylsphingosine in GBA1 Mutation Carriers with and without Parkinson’s Disease. Mov. Disord. 37, 416–421 (2022).

69. Blumenreich, S. et al. Elevation of gangliosides in four brain regions from Parkinson’s disease patients with a GBA mutation. npj Park. Dis. 8, (2022).

70. Lerche, S. et al. The Mutation Matters: CSF Profiles of GCase, Sphingolipids, α-Synuclein in PDGBA. Mov. Disord. 36, 1216–1228 (2021).

71. Brody, E. M. et al. GPNMB Biomarker Levels in GBA1 Carriers with Lewy Body Disorders. Mov. Disord. 39, 1065–1070 (2024).

72. Roberts, B. et al. Systematic gene tagging using CRISPR/Cas9 in human stem cells to illuminate cell organization. Mol. Biol. Cell (2017) doi:10.1091/mbc.E17-03-0209.

73. Roberts, B. et al. Fluorescent Gene Tagging of Transcriptionally Silent Genes in hiPSCs. Stem Cell Reports 12, 1145–1158 (2019).

74. Chen, Y., et al. *A versatile polypharmacology platform promotes cytoprotection and viability of human pluripotent and differentiated cells*. Nature Methods vol. 18 (Springer US, 2021).

75. Kunkle, B. W. et al. Genetic meta-analysis of diagnosed Alzheimer’s disease identifies new risk loci and implicates Aβ, tau, immunity and lipid processing. Nat. Genet. 51, 414–430 (2019).

76. Nalls, M. A. et al. Identification of novel risk loci, causal insights, and heritable risk for Parkinson’s disease: a meta-analysis of genome-wide association studies. Lancet Neurol. 18, 1091–1102 (2019).

77. Fiume, M. et al. CRISPResso2 provides accurate and rapid genome editing sequence analysis. Nat. Biotechnol. 37, 220–224 (2019).

78. Williams, D., et al. High-throughput screening for small-molecule stabilizers of misfolded glucocerebrosidase in Gaucher disease and Parkinson’s disease. Proc. Natl. Acad. Sci. 121, e2406009121 (2024).

79. Deen, M. et al. A versatile fluorescence-quenched substrate for quantitative measurement of glucocerebrosidase activity within live cells. PNAS 119, 1–10 (2022).

80. Bielawski, J. et al. Sphingolipid Analysis by High Performance Liquid Chromatography-Tandem Mass Spectrometry (HPLC-MS/MS). in Sphingolipids as Signaling and Regulatory Molecules (eds. Chalfant, C. & Poeta, M. Del) 46–59 (Springer New York, 2010). doi:10.1007/978-1-4419-6741-1_3.

81. Bielawski, J. et al. Comprehensive Quantitative Analysis of Bioactive Sphingolipids by High-Performance Liquid Chromatography--Tandem Mass Spectrometry. in Lipidomics: Volume 1: Methods and Protocols (ed. Armstrong, D.) 443–467 (Humana Press, 2009). doi:10.1007/978-1-60761-322-0_22.

82. Bligh, E. G. & Dyer, W. J. A RAPID METHOD OF TOTAL LIPID EXTRACTION AND PURIFICATION. Can. J. Biochem. Physiol. 37, 911–917 (1959).

