## Supplementary Figures for "Bidirectional regulation of glycoprotein nonmetastatic melanoma protein B by β-glucocerebrosidase deficiency in *GBA1* isogenic dopaminergic neurons from a patient with Gaucher disease and parkinsonism"

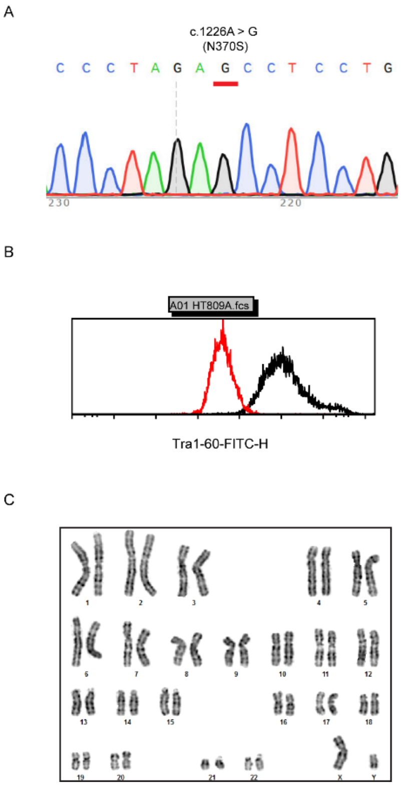


**Figure S1: Characterization of the patient-derived iPSC line HT809.**

**A.** Homozygous *GBA1* N370S variants in HT809 confirmed by Sanger sequencing. **B.** The expression of TRA-1-60, a human pluripotent stem cell marker, in HT809 iPSCs. **C**. Normal karyotype of HT809.

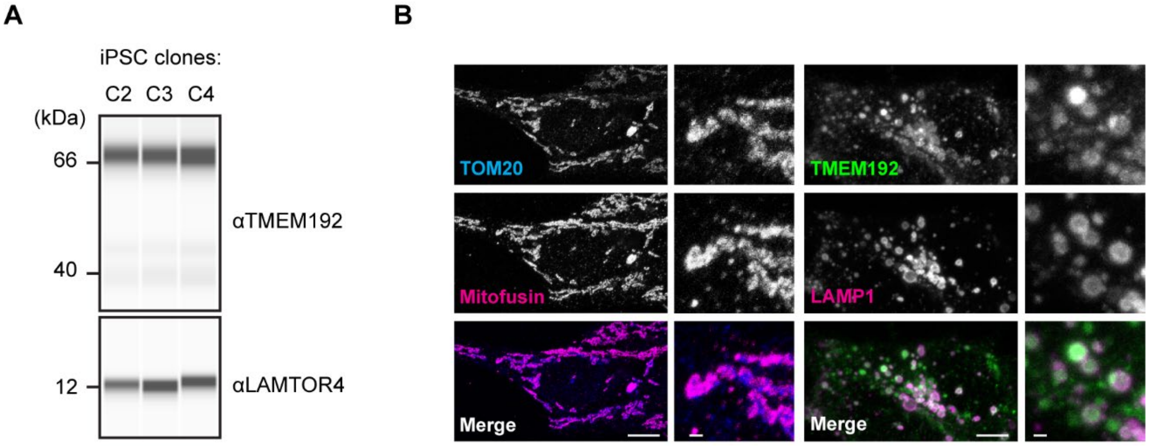


**Figure S2: The expression of TMEM192-GFP-3xHA in HT809.**

**A**. TMEM192-GFP-3xHA expression in edited HT809 iPSC clones. **B.** Localization of TOM20-tagBFP-2xStrep to mitochondria and Tmem192-GFP-3xHA to lysosomes. Scale bar, 10 μm; insert scale bar, 1 μm.

A

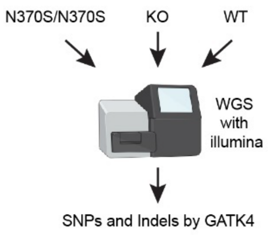


| #1 N370S/N370S | #2 KO | #3 WT |
|---|---|---|
| 155205634 GBA | 155205634 GBA | |
| 171076966 FMO3 | 171076966 FMO3 | 171076966 FMO3 |
| 171083242 FMO3 | 171083242 FMO3 | 171083242 FMO3 |
| 179545050 NPHS2 | 179545050 NPHS2 | 179545050 NPHS2 |
| 60720246 BCL11A | 60720246 BCL11A | 60720246 BCL11A |
| 172305177 DCAF17 | 172305177 DCAF17 | 172305177 DCAF17 |
| 227892720 COL4A4 | 227892720 COL4A4 | 227892720 COL4A4 |
| 151936677 CCDC170 | 151936677 CCDC170 | 151936677 CCDC170 |
| 151948366 CCDC170;ESR1 | 151948366 CCDC170;ESR1 | 151948366 CCDC170;ESR1 |
| 97367834 FBP1 | 97367834 FBP1 | 97367834 FBP1 |
| 17298125 NUCB2 | 17298125 NUCB2 | 17298125 NUCB2 |
| 52508989 ATP7B | 52508989 ATP7B | 52508989 ATP7B |
| 13007458 GCDH | 13007458 GCDH | 13007458 GCDH |
| 13010520 GCDH | 13010520 GCDH | 13010520 GCDH |
| 13010643 GCDH | 13010643 GCDH | 13010643 GCDH |
| 49469087 FTL | 49469087 FTL | 49469087 FTL |
| 15326 NONE;NONE | 15326 NONE;NONE | 15326 NONE;NONE |

B

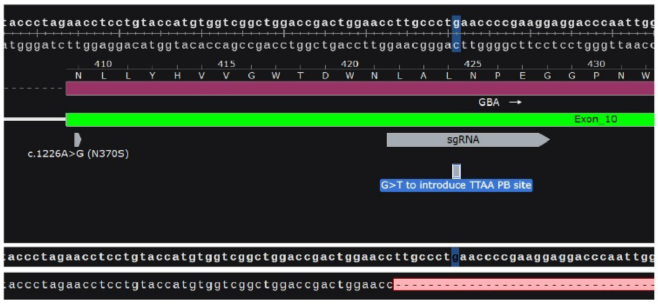


C

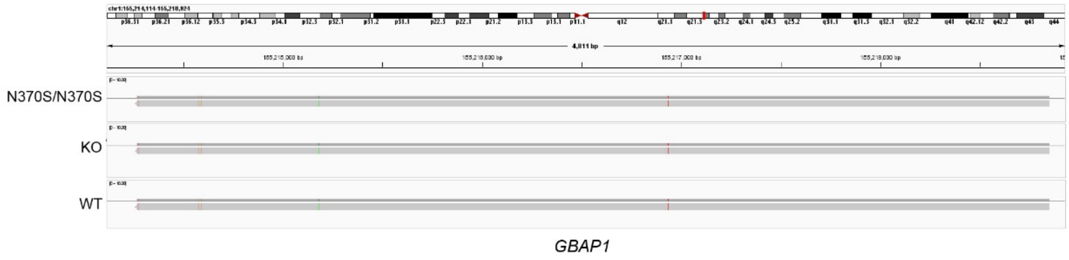


**Figure S3: Specific editing of *GBA1* in HT809.**

**A.** Variants with a gnomAD allele frequency (AF) of <0.001 and a combined annotation-dependent depletion (CADD) phred score of >30 in HT809 *GBA1* isogenic iPSC lines. Note the absence of *GBA1* N370S variants in the WT line. **B.** gRNA designed to specifically target *GBA1* near the N370S site. **C.** No changes were introduced to *GBAP1* in the isogenic lines as demonstrated by PacBio HiFi long-read amplicon sequencing.

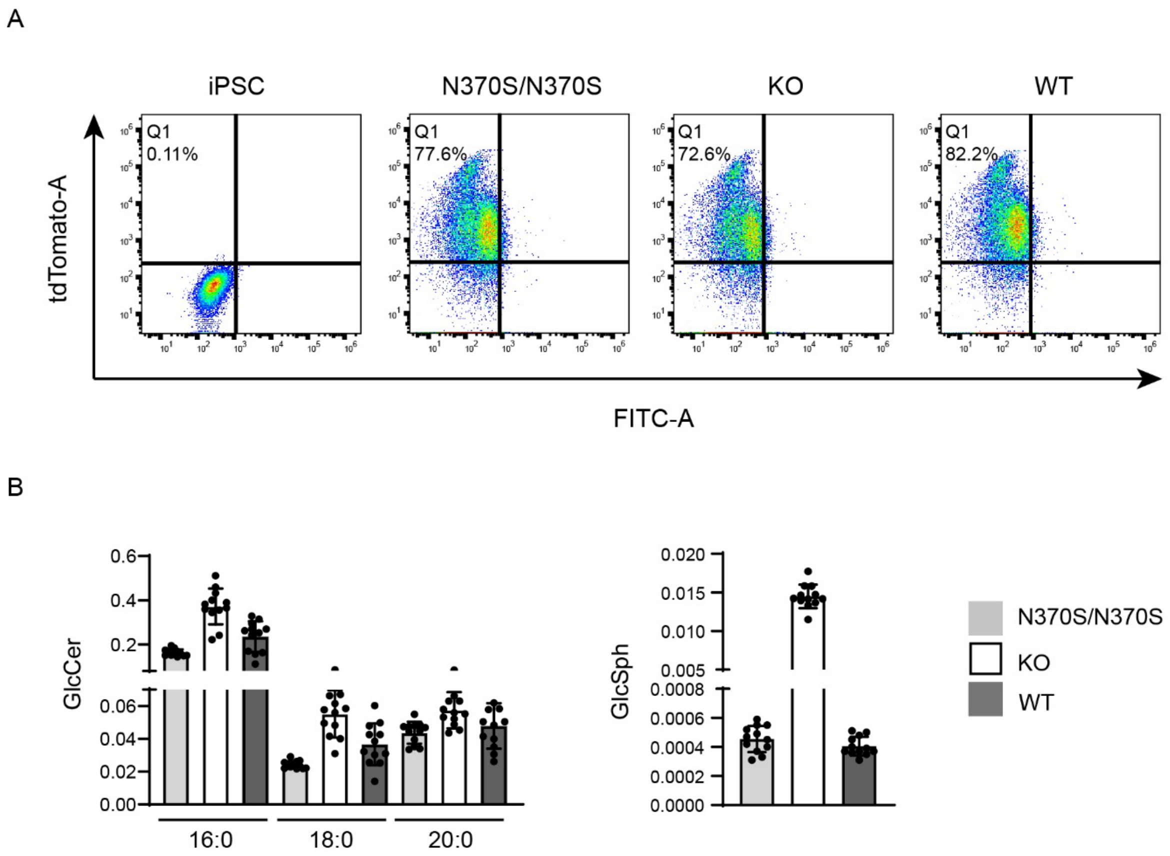


**Figure S4: HT809 isogenic iPSC DAN differentiation efficiency and GlcCer and GlcSph levels.**
**A.** DAN differentiation efficiency of WT, N370S/N370S, and KO iPSCs quantified based on tdTomato+ cell population on day 25 of differentiation. **B.** GlcCer and GlcSph quantification in WT, N370S/N370S, and KO iPSCs. Data was normalized with cell counts.

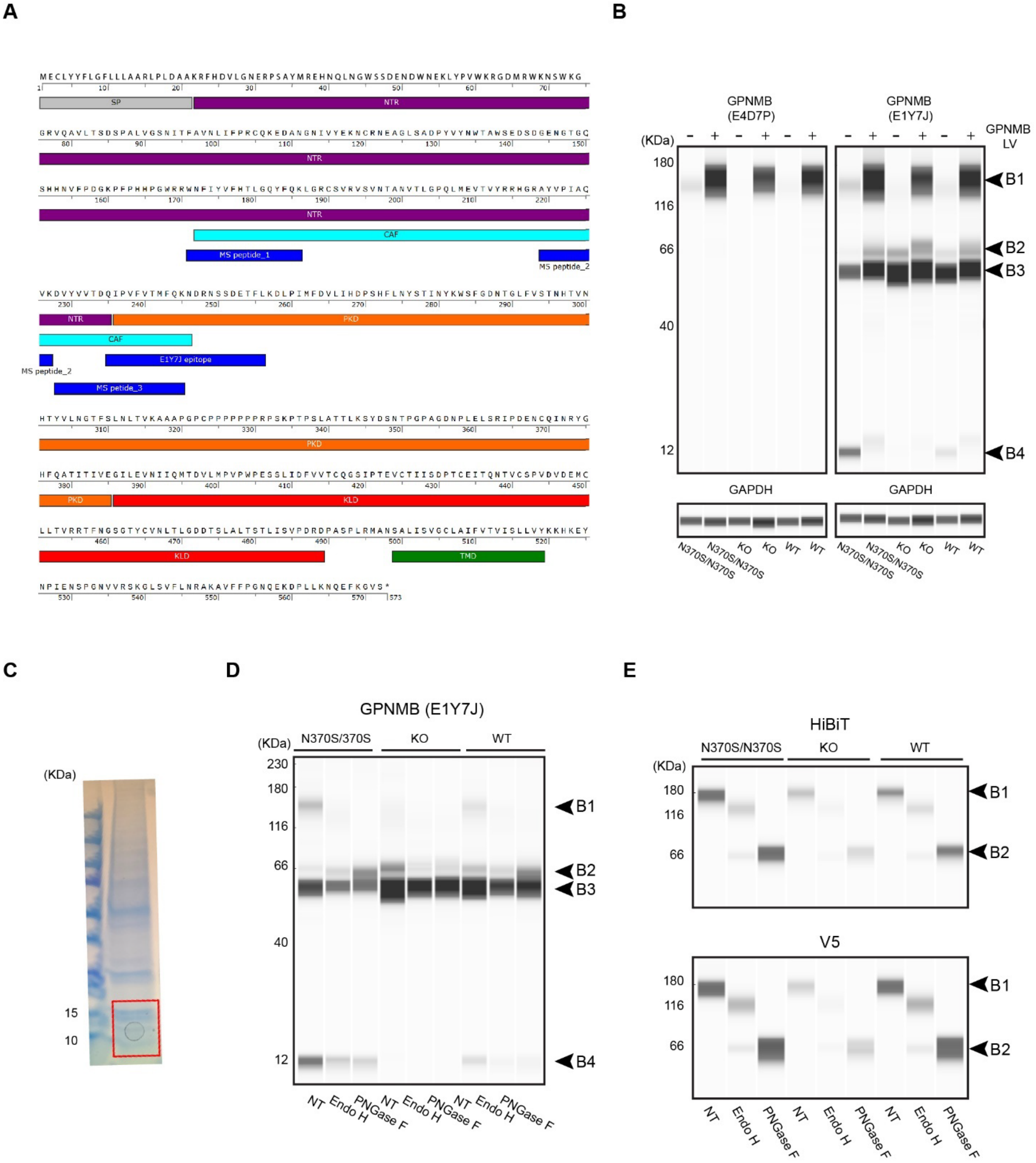


**Figure S5: GPNMB in DANs.**

A, GPNMB protein sequence, domain structure, and mapping of MS. peptides and antibody epitopes. B, Expression of HiBiT-GPNMB-V5 in transduced DANs. C, Proteins extracted from DAN lysosomes were separated and gel slice between 10 and 15 kDa were analyzed by mass spectrometry for protein identification. D.E. GPNMB glycosylation analysis in DANs using E1Y7J, HiBiT, and V5 antibodies.

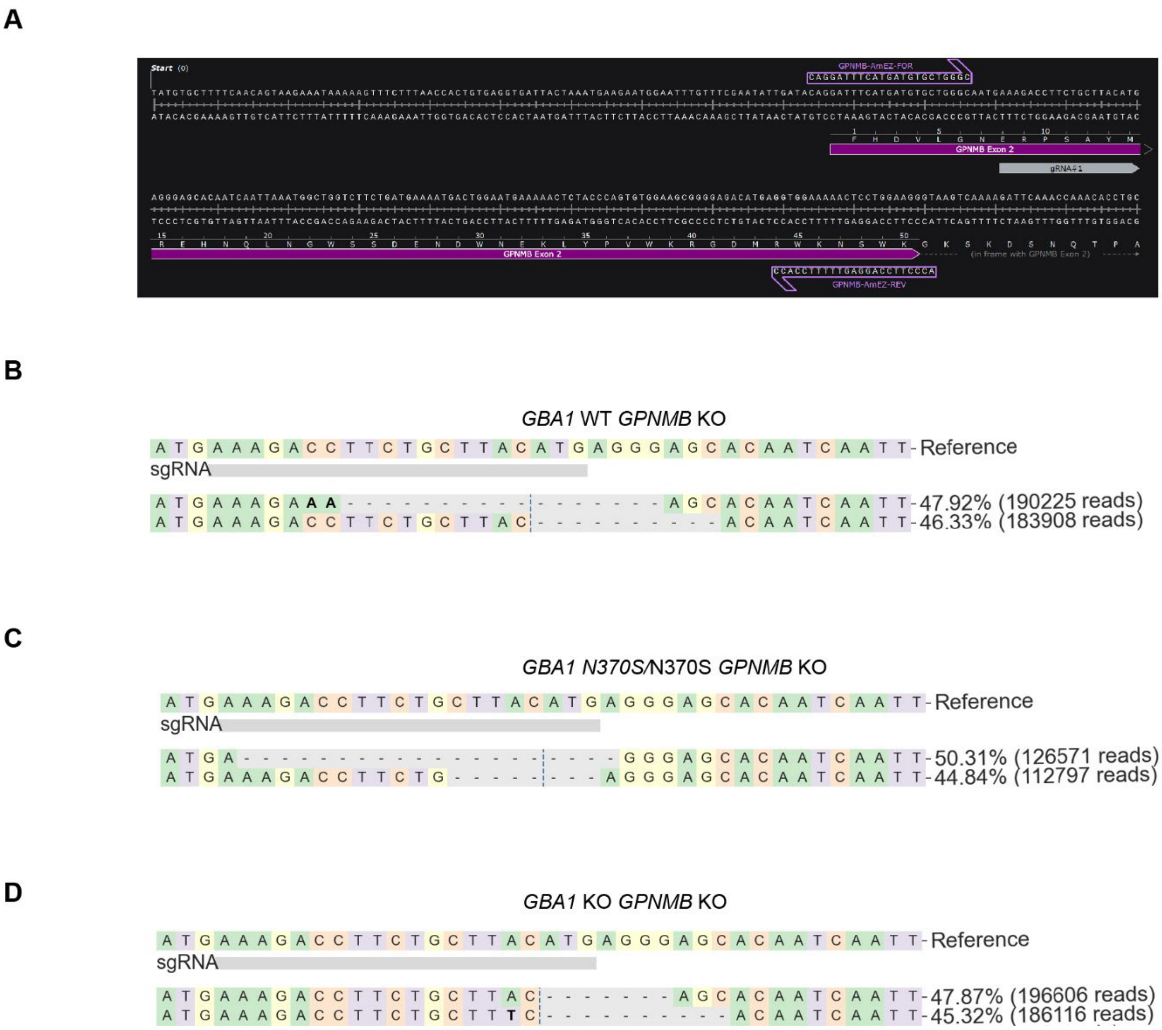


**Figure S6: GPNMB knockout in HT809 *GBA1* isogenic iPSCs.**

**A.** gRNA designed to target *GPNMB* exon 2. **B,C,E,** short-read amplicon sequencing confirming INDEL mutations in both *GPNMB* alleles in all HT809 *GBA1* isogenic iPSCs.
